# Characterization of Fetal Cortical Development Using Spectral Analysis of Gyrification (SPANGY)

**DOI:** 10.64898/2026.07.14.736987

**Authors:** Harvey Dienye, Angeline Mihailov, Thomas Sanchez, Gerard Martí-Juan, Raquel G López, Léo Pomar, Joanna Sichitiu, Vincent Dunet, Mériam Koob, Elisenda Eixarch, Aurelie Manchon, Nadine Girard, Mathieu Milh, David Germanaud, Miguel A. Gonzalez Ballester, Oscar Camara, Gemma Piella, Meritxell Bach Cuadra, François Rousseau, Julien Lefèvre, Olivier Coulon, Guillaume Auzias

## Abstract

The prenatal period of human brain development is critical for mental health and cognition across the entire lifespan. During this period, the cortex undergoes a dramatic transformation from a smooth lissencephalic surface into an elaborately folded structure, a process whose precise characterization is essential for understanding neurodevelopmental trajectories. This study represents the first application of Spectral Analysis of Gyrification (SPANGY) to a large multi-centric fetal brain MRI dataset (635 subjects, 20–38 weeks gestational age). SPANGY characterizes geometric variations on a surface based on the wavelength of folds, hence, providing a quantitative local description of gyrification at the individual level. Using rigorous normative modeling (GAMLSS) and statistical harmonization (ComBat-GAM), we established age-specific reference trajectories for multi-scale gyrification features (spectral frequency bands). We provide the first ever quantification of the temporally-ordered emergence of cortical folding in successive waves during the entire second half of pregnancy: the earliest-emerging low frequency, deep fissures are progressively superseded by the accelerating expansion of higher frequency folds. The normative curves provide the first step in taking prenatal neurodevelopmental assessment from qualitative inspection into a rigorous statistical inference, creating an objective reference against which deviations from healthy brain growth could be caught earlier.

## 1. Introduction

The prenatal period of human brain development is critical for mental health and cognition across the entire lifespan (Walhovd et al., 2024). During this time, an ensemble of complex and dynamic biological processes are orchestrated, in parallel with a dramatic increase of cortical surface area resulting in cortical folding.

Recent advances in cellular neurobiology have comprehensively described the spatio-temporal organization of the fetal brain at the cellular level, highlighting the role and timing of intricate biological processes such as neural progenitor proliferation and neuronal migration, myelination and synaptogenesis (Kostović et al., 2019; Llinares-Benadero and Borrell, 2019).

At the macroscopic level, fetal brain magnetic resonance imaging (MRI) has advanced significantly through the integration of artificial intelligence (AI) (Sanchez et al., 2025; Uus et al., 2023; Xu et al., 2023), allowing us to study cortical folding in vivo during the critical prenatal period. In contrast to postnatal imaging, fetal MRI enables the direct observation of gyrification as it progresses and captures the transformation of the cortex from a rather smooth lissencephalic cortex to an elaborately folded mature configuration (Ciceri et al., 2024; Gholipour et al., 2017). Quantitative characterisations of this trajectory, based on local tissue expansion (Rajagopalan et al., 2011), curvature- and contour-based gyrification indices on routine 2D acquisitions (Yehuda et al., 2023), and perinatal gyrification analysis (Mihailov et al., 2025), have delineated the spatiotemporal sequence of cortical folding and gyrification, and revealed regional asymmetries and sex-specific growth trajectories (Studholme et al., 2020).

The microscopic scale of cellular processes occurring at resolutions far below current in vivo imaging capabilities, creates a fundamental gap between cellular understanding and practical measurement. This disconnect necessitates complementary approaches that can bridge scales, providing quantitative measures of folding patterns that both emerge from and complement their cellular origins. Addressing this knowledge gap is particularly challenging since the measures of the evolution in time are restricted to the macroscopic scale (typically on the order of mm for in-vivo neuroimaging such as MRI), while the access to microscopic scale requires histology and is thus not compatible with the characterization of the dynamics of brain maturation. The precise assessment of spatial and temporal dimensions are thus mostly independent due to technical limitations in the available recording approaches, and the integration of time and space relies on either correlational studies or modeling.

For correlational studies, one the most advanced investigations combined in-vivo MRI with (post-mortem) histology in non-human primate to document the synergistic interactions between brain maturation at the cellular level and the dynamics of cortical folding (Wang et al., 2017). Another approach consists in characterizing abnormal cortical folding patterns in some neurodevelopmental disorders for which the pathophysiology is sufficiently documented, to push the interpretation of the underlying biological processes, such as in Williams syndrome for which the genetic pathways are known (Kippenhan et al., 2005; Thompson et al., 2005), or corpus callosum agenesis (Schwartz et al., 2021) and polymicrogyria (Lamballais et al., 2020). In combination with the aforementioned MRI studies on healthy fetuses, this literature refined our understanding of the dynamics of early brain development. It clarified the origin of the correlations between cortical geometry and cellular organization, observed in human adults more than one century ago (Economo et al., 2008) and confirmed with advanced techniques recently (Wagstyl et al., 2018).

On the other hand, biomechanical modeling represents a prominent approach to understanding the origin and emergence of cortical folding during fetal development. These models conceptualize the developing brain as a multi-layered structure comprising an outer cortical layer, inner white matter core, and intermediate proliferative zones, where differential growth rates between layers drive mechanical instabilities that manifest as cortical folds (Lefèvre and Mangin, 2010; Yin et al., 2025). The fundamental principle posits that faster tangential expansion of the cortical plate relative to underlying structures generates compressive stresses that are relieved through buckling, producing the characteristic convoluted morphology of the human brain (Garcia et al., 2018; Tallinen et al., 2016). While computational implementations of these models have achieved some success in reproducing folding patterns and demonstrating how variations in growth parameters, tissue properties, and initial geometry influence gyrification complexity (Wang et al., 2021), they face substantial limitations for practical application. The complexity of high-resolution biomechanical simulations, requiring extensive parameter tuning and validation, as well as a better understanding of the many mechanisms interacting during fetal development, further limit their practical utility.

In this context, our work investigates the multi-scale nature of cortical morphogenesis, using an original approach grounded in explicit geometric assumptions that provide complementary interpretations and perspectives compared to classical MRI-derived features.

Intuitively, global metrics like gyrification index produce a single value per hemisphere (Mihailov et al., 2025; Rajagopalan et al., 2011; Yehuda et al., 2023), which lacks the granularity necessary for a detailed understanding of the evolution of cortical folding. To address this limitation, alternative approaches have focused on metrics computed at the scale of lobes, regions or sulci. Sulcal depth measurements have been employed to track the emergence and deepening of individual cortical folds throughout gestation, providing spatially detailed characterization of primary sulcal development (Solhtalab et al., 2025; Xu et al., 2022). Sulcal identification and morphometry approaches have demonstrated that individual sulcal patterns capture clinically relevant developmental variations and can distinguish between typical and atypical folding trajectories (De Vareilles et al., 2023). Surface-based measures including mean curvature, Gaussian curvature, and sulcal pit depth have also been utilized to characterize local geometric properties of the developing cortical surface (Bartha-Doering et al., 2023; Clouchoux et al., 2012; Demirci and Holland, 2022; Im and Grant, 2019; Yun et al., 2020). These local metrics offer improved spatial specificity compared to global indices, but they rely on regions or sulci defined a priori, that have to be labelled on each individual brain. The labeling is often propagated from an atlas using registration techniques such as ANTS (Tustison et al., 2021) or MSM (Robinson et al., 2018). Distortions and/or inaccuracies induced by the registration can thus affect the measures and limit the scale of observation to relatively large regions and most prominent sulci, potentially overlooking the more subtle and variable folding patterns that emerge progressively throughout the second and third trimesters.

In this work, we address these limitations by presenting the first application of Spectral Analysis of Gyrification (SPANGY) (Germanaud et al., 2012) on a large multicentric fetal neuroimaging dataset. As detailed below, SPANGY has been designed to quantitatively analyse folding patterns based on the spectral decomposition of the mean curvature, computed at the individual level, thus bypassing the registration step. The rich, multi-scale information extracted by SPANGY enables, for the first time, the construction of normative developmental models that characterize how each SPANGY frequency band evolves throughout pregnancy. We establish the first normative age-specific reference trajectories for cortical folding during the fetal period.

Furthermore, we demonstrate that the developmental trajectories of SPANGY-derived frequency bands provide quantitative support of the idea that the cortex morphology develops in successive waves of folding. We document the hierarchical, temporally-ordered emergence of cortical sulci during fetal brain development, hereby providing the first quantitative confirmation of previously reported qualitative observations (Chi et al., 1977; De Vareilles et al., 2023).

In addition, the integration of SPANGY with normative modeling is essential for understanding the heterogeneity of neurodevelopmental trajectories and for identifying subtle anomalies that may presage later neurodevelopmental disorders (Bethlehem et al., 2022). These scale-specific normative references represent a promising move from descriptive characterization to predictive quantification in prenatal neurodevelopmental assessment, with potential applications in early detection and monitoring of atypical brain development.

## 2. Materials and Methods

### 2.1. Data

To effectively and reliably characterize the dynamics of cortical development, large samples are essential. To this regard, our study was run on the same datasets as in Sanchez et al., 2026, which includes 4 datasets obtained from 4 different sites (MarsFet, Fetal dHCP, BCNatal and CHUV), combining research-dedicated acquisitions with retrospective access to data acquired in clinical routine. Exclusion criteria applied to all datasets were designed through iterative consensus meetings among a multidisciplinary clinical team, incorporating evidence-based guidelines from current literature and established clinical protocols. These criteria encompassed multiple domains including maternal medical history, pregnancy complications, and fetal anomalies detected through prenatal screening. Specifically, cases were excluded if they presented with:

- Documented chromosomal abnormalities or genetic syndromes
- Structural brain malformations identified through conventional neuroimaging (prenatal ultrasound and prenatal MRI)
- Maternal conditions known to impact fetal neurodevelopment (e.g., diabetes mellitus, substance abuse, teratogenic medication exposure)
- Pregnancy complications such as intrauterine growth restriction or oligohydramnios
- Evidence of fetal distress or abnormal biophysical profiles during the assessment period
- No consent from the expecting parents.

Applying these criteria led to a combined cohort of 635 subjects. A basic description of each dataset can be seen in Table 1. A detailed description of each dataset is available in Sanchez et al., 2026.

**Table 1.** Overview of the multicentric fetal neuroimaging dataset. The cohort includes data from four independent centers, detailing the sample size (n), biological sex distribution, magnetic resonance imaging field strength, and gestational age characteristics.

| Dataset | *n* | Sex Ratio (F:M:unknown) | Scanner (3T:1.5T) | Age Range (Weeks of Gestational Age) |
| --- | --- | --- | --- | --- |
| MarsFet | 216 | 83:90:43 | 114:102 | 20.85 – 38.00 |
| dHCP | 292 | 133:157:2 | 292:0 | 20.86 – 38.29 |
| BCNatal | 91 | 33:55:3 | 59:32 | 26.43 – 38.86 |
| CHUV | 36 | 12:24:0 | 3:33 | 21.00 – 35.71 |

### 2.2. 3D image reconstruction, tissue segmentation and surface extraction using Fetpype

In this study, we implemented the same image processing pipeline as detailed in Sanchez et al., 2026. Standardized processing across all datasets was achieved using Fetpype, a publicly available computational framework developed for consistent preprocessing, reconstruction, and segmentation of fetal brain MRI data (Sanchez et al., 2025). Fetpype is publicly available on Github: https://github.com/fetpype/fetpype.

### 2.3. Assessment of the quality of the surfaces

Poor image quality is expected to impact all steps of the image processing pipeline described above. In this work, we paid particular attention to the quality of the surfaces that are the input of the SPANGY analysis.

#### Mesh Quality Assessment

Quality control was performed through visual inspection of all 1,270 cortical surface meshes (635 subjects × 2 hemispheres), with each mesh assigned a quality rating on a 5-point ordinal scale. Meshes with severe reconstruction failures were assigned low scores based on fundamental anatomical criteria. A score of 1 indicated complete reconstruction failure, where the generated surface bore no recognizable resemblance to cerebral cortical anatomy. A score of 2 was assigned to meshes that demonstrated a generally brain-like morphology but exhibited major anatomical inconsistencies, such as missing lobes, grossly distorted sulcal-gyral patterns, or substantial topological errors that precluded meaningful anatomical interpretation.

Meshes receiving scores of 3 or higher demonstrated correct cortical morphology and anatomical consistency. Within this range, scores were differentiated based on the severity and extent of surface-level artifacts. These artifacts included:

1. **Surface noise**: Visible irregularities, roughness, or high-frequency oscillations on the cortical surface that deviated from the expected smooth geometry
2. **Segmentation artifacts**: Reconstruction errors stemming from possible inaccuracies in the underlying tissue segmentation, such as erroneous inclusion of non-cortical tissue, or locally irregular surface geometry.

Higher scores (4-5) were reserved for meshes with minimal surface noise and few segmentation artifacts, while a score of 3 indicated acceptable anatomical structure but noticeable surface quality issues that, while not compromising overall morphology, were visually apparent upon inspection.

Surfaces with a score below 3 were excluded from the analysis. As a result, the final sample of surfaces (including both left and right hemispheres) retained for the SPANGY analysis comprised 312 surfaces from MarsFet, 585 from dHCP, 108 from BCNatal, and 50 from CHUV.

### 2.4. SPANGY Application

Spectral Analysis of Gyrification (SPANGY) is a framework developed by Germanaud et al. (Germanaud et al., 2012) to characterize cortical folding through spectral decomposition of the cortical surface curvature. The technique operates by computing eigenfunctions of the Laplace-Beltrami operator on the cortical mesh surface that serve as a basis to decompose the mean curvature field. These eigenfunctions are then grouped into frequency bands following a frequency-doubling principle, where each successive band contains eigenfunctions with approximately twice the spatial frequency of the preceding band, starting with the first eigenfunction. These bands capture the hierarchical organization of cortical folding, from broad, low-frequency undulations representing global brain shape to fine, high-frequency features corresponding to local geometric variations (Germanaud et al., 2012).

As illustrated on Fig. 1, the spectral decomposition in SPANGY generates seven distinct frequency bands, labeled B0 through B6, each representing progressively finer spatial scales of cortical surface patterns. Understanding these frequency bands is essential to interpreting SPANGY’s output (Germanaud et al., 2012). While the lower bands capture global hemispheric geometry, patterns indicative of true cortical folding typically emerge at band B4. The most substantial contributions to folding power are concentrated in bands B4, B5, and B6, which correspond to the hierarchical sulcal and gyral features that constitute cortical gyrification (Dubois et al., 2019; Germanaud et al., 2012). For this reason, our analysis focused on bands B4, B5, and B6, as they capture the biologically relevant scales of cortical maturation for this study.

**Fig. 1.**
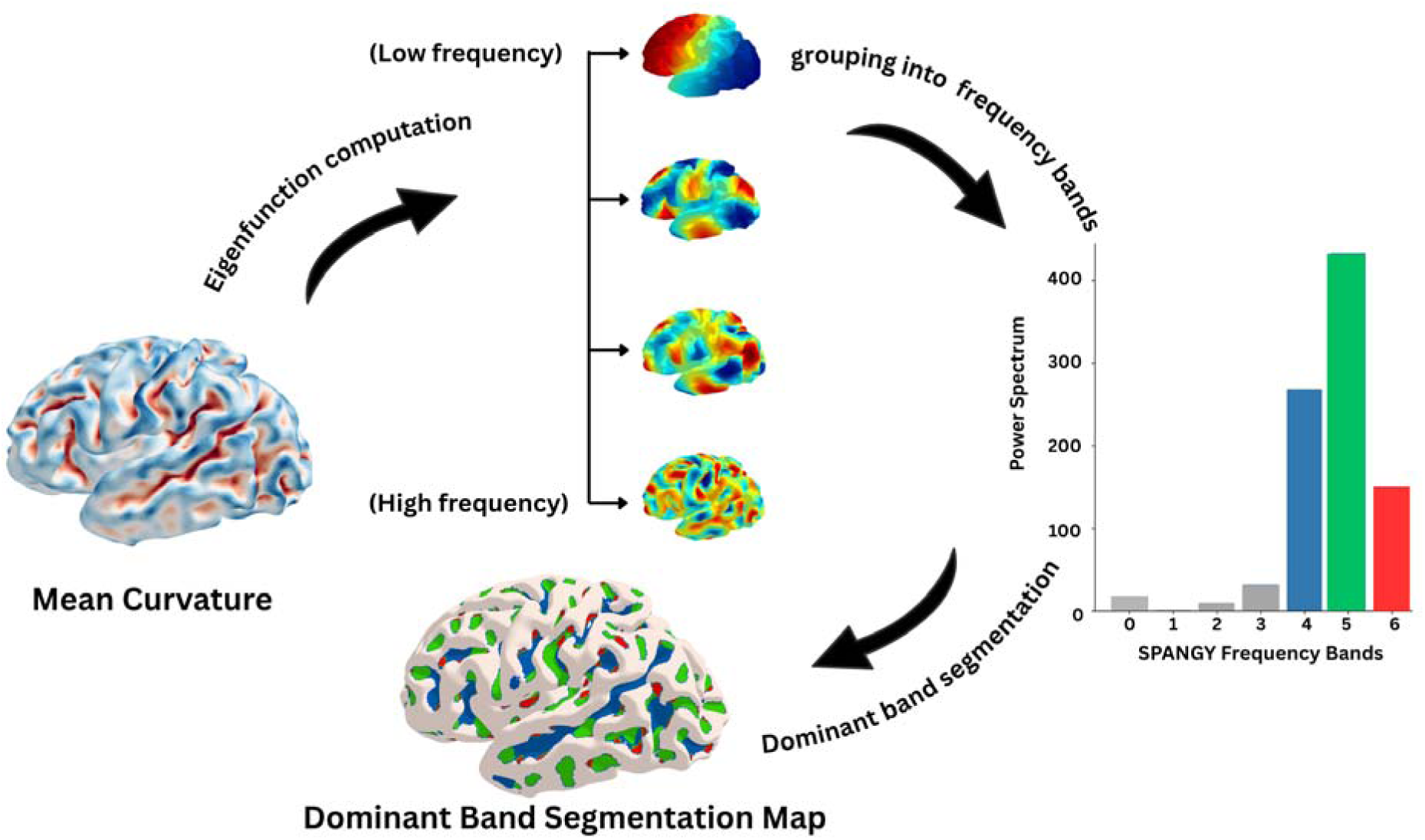
Overview of the SPANGY methodology. The input consists of the mean curvature mapped onto the reconstructed fetal cortical mesh (left). Spectral decomposition is performed by computing the eigenfunctions of the Laplace-Beltrami operator, which provide a natural basis for the surface. These eigenfunctions are grouped into successive frequency bands via a frequency-doubling principle, capturing a hierarchy of spatial scales. The final dominant frequency segmentation identifies the specific scale that locally characterizes the cortical folding.

Following spectral decomposition, we performed segmentation of the cortical surface based on the dominant frequency band at each vertex. For each vertex on the cortical mesh, SPANGY quantifies the power contribution from each frequency band. The dominant band at a given vertex is defined as the frequency band that contributes the maximum spectral power at that location (Germanaud et al., 2012). By assigning each vertex to its dominant frequency band, the entire cortical surface is partitioned into mutually exclusive regions, each characterized by a specific spatial frequency.

As in Germanaud et al., 2012, the visualization of the dominant frequency band segmentation was restricted to the sulcal regions primarily to maintain the clarity of the resulting maps allowing for a focused and representative assessment of folding complexity without redundancy.

To obtain a global measure of cortical folding complexity, we calculated the total folding power for each subject by summing the spectral power values across all seven frequency bands (B0 through B6). This metric provides a comprehensive index of overall gyrification that integrates information across all spatial scales, functioning analogously to a spectral gyrification index (Rabiei et al., 2017).

In addition to the global measure, we extracted band-specific power values by isolating the spectral contributions for bands B4, B5, and B6. These band-specific power measures quantify the absolute contribution of each folding band to the overall cortical architecture and enable tracking of how different folding bands evolve across development.

To characterize the spatial extent of different folding patterns, we calculated the percentage of cortical surface area dominated by each band (B4, B5, and B6). This was computed by dividing the surface area of vertices assigned to each band by the total cortical surface area.

We also computed the relative band power for B4, B5, and B6 by normalizing each band’s power against the total folding power to capture the proportional contribution of each folding level to the overall spectral composition, providing a size-invariant characterization that remains independent of absolute brain volume or total folding magnitude.

All extracted metrics (including total folding power, band-specific powers (B4, B5, B6), percentage surface areas for each band, and relative band powers) were systematically compiled for each subject and hemisphere. These quantitative features constituted the input variables for subsequent normative modeling analyses.

The full SPANGY code is available on GitHub: https://github.com/brain-slam/slam, and https://github.com/fetpype/Spangy-Fet.

### 2.5. Adaptation of SPANGY to fetal data

#### Tuning of SPANGY Parameters for Fetal Brain Analysis

SPANGY was originally developed and validated for the analysis of adult cortical surfaces (Germanaud et al., 2012), necessitating careful optimization of methodological parameters for application to fetal brain data. Fetal neuroimaging presents unique technical challenges compared to adult or pediatric studies, and our multi-site acquisition protocol introduced additional sources of variance requiring systematic parameter evaluation. We assessed and optimized two critical parameters (surface smoothing and number of eigenfunctions) to ensure accurate spectral decomposition of fetal cortical surfaces.

#### Surface Smoothing

SPANGY depends on the regularity of the mean curvature field. Mean curvature, being a second-order geometric property, exhibits high sensitivity to local surface perturbations and noise (Pienaar et al., 2008). Random high-frequency components introduced by mesh irregularities can contaminate the spectral decomposition and generate artifactual contributions to higher frequency bands, potentially confounding genuine folding patterns with technical noise.

To address this issue, we applied iterative Laplacian smoothing to all cortical surface meshes prior to SPANGY analysis. The algorithm updates each vertex position based on the weighted average of its neighboring vertices, effectively applying a low-pass filter to the surface coordinates. We systematically evaluated it through 5, 10, and 20 iterations to determine the optimal balance between noise reduction and feature preservation. Through visual inspection and quantitative assessment of mesh quality metrics (including vertex regularity and surface curvature distributions), we determined that 5 iterations provided sufficient smoothing to eliminate acquisition artifacts while preserving critical sulcal features necessary for SPANGY analysis. This smoothing level effectively attenuated mesh irregularities and noise-related high-frequency components while maintaining the anatomical fidelity of sulcal and gyral features. An illustration of these steps can be seen in the supp. Fig. 1.

#### Number of Eigenfunctions (N)

The spectral decomposition in SPANGY requires specification of the number of eigenfunctions computed from the Laplace-Beltrami operator. This parameter determines the maximum spatial frequency captured in the analysis and must be sufficient to represent the full range of folding patterns present in the data (Germanaud et al., 2012; Lefèvre et al., 2018). An insufficient number of eigenfunctions truncates the frequency spectrum prematurely, leading to incomplete characterization of finer folding patterns, while an excessive number of eigenfunctions increases computational cost without providing additional information once the mesh’s intrinsic frequency content has been fully captured.

We computed 5000 eigenfunctions for each hemispheric surface. Inspection of eigenvalue spectra and visual assessment of high-order eigenfunctions across the gestational age range confirmed that this number adequately captured all meaningful spatial frequencies present in fetal cortical surfaces, with the higher-order eigenfunctions showing negligible contribution to cortical folding power, indicating that the frequency spectrum had been fully sampled. This is illustrated on Supp. Fig. 2.

### 2.6. Normative Modelling

Normative modeling is a statistical framework that characterizes typical patterns of brain development by mapping the expected trajectory of neuroimaging features as a function of age or other covariates (Marquand et al., 2016). This approach has become increasingly important in neuroscience as it enables the quantification of individual deviations from normative trajectories, facilitating the identification of atypical developmental patterns and potential biomarkers for neurodevelopmental disorders (Bethlehem et al., 2022; Rutherford et al., 2023).

Since this study is the first application of SPANGY to data acquired from multiple sites, we first conducted a systematic evaluation of site-related variance. Across all extracted features, including a random effect for the site was seen to be significant in achieving a better fit for the models. The p-values of the effect of the site on the different frequency bands are reported in the supplementary materials (Supp.Table 1). The significance of the random effects specification demonstrated that site-level heterogeneity can affect optimal model fit, thereby establishing the empirical rationale for our subsequent harmonization approach.

Following this evaluation, we performed statistical harmonization using ComBat-GAM to remove site-related effects while preserving biological variability (Pomponio et al., 2020). We used the dHCP dataset as the reference site, ensuring that scanner and acquisition-related effects were mitigated across the multi-site cohort.

After harmonization, we implemented Monte Carlo Cross-Validation (MCCV) on the training data, to ensure robust model estimation and evaluation. We randomly assigned 80% of the data to the training set and 20% to the test set. We performed 20 rounds of MCCV with stratified sampling with respect to gestational age and subject, i;e we also ensured that both hemispheres of a single subject fell into the same group (test or training), to avoid introducing bias in our models. This approach allowed us to assess model performance across different data partitions and mitigate the risk of overfitting. We systematically investigated different normative models of increasing complexity to identify the optimal framework for modeling our cortical gyrification data.

As proposed in Dinga et al., 2021, the model selection process followed a hierarchical approach, beginning with simpler models and progressively incorporating additional complexity. We started with a linear model, then progressed to a Generalized Additive Model (GAM) with smooth modeling of the mean, followed by a GAM incorporating smooth modeling of both mean and variance. We then implemented a GAMLSS model (Stasinopoulos and Rigby, 2012) using the Sinh-Arcsinh (SHASH) distribution with smooth modeling of mean and variance, while kurtosis and skewness were modeled as fixed values. Finally, we evaluated a full GAMLSS SHASH model with smooth modeling of all distribution parameters (mean, variance, skewness, and kurtosis).

Model selection was based on the log score evaluated on our test set, which quantifies the predictive performance of each model on unseen data (Dinga et al., 2021). This criterion prioritizes models that generalize well beyond the training data, balancing goodness of fit against the risk of overfitting through out-of-sample validation. This model selection procedure was applied independently for each morphometric feature. Following cross-validation and model selection, we generated normative developmental curves for each feature using the best-performing model. These curves characterize the trajectories of cortical folding across gestational age and provide the reference framework for subsequent analyses.

## 3. Results

In this section we report and describe the normative models fitted to the features derived from the SPANGY analysis, reserving their biological interpretation for the Discussion. We present normative curves for nine features, corresponding to three spectral metrics computed for each of the three highest-frequency bands (B4-B6). The first, absolute band power, captures how the spectral power of each band evolves over gestation. The second, band surface area percentage, describes how the proportion of cortical surface occupied by each band changes, offering a surface-level view of the emerging folding pattern. The third, relative band power, expresses each band’s power as a fraction of the total; this normalisation facilitates direct comparison with earlier SPANGY studies (Dubois et al., 2019).

### 3.1. Normative curves

The model selection results indicated that more sophisticated models were generally preferred over basic linear approaches. Details regarding the selected model for each morphometric feature, including mean log scores from the 20 MCCV rounds, are provided in Supp. Table 2.

Upon completing model development, selection and optimization, we fitted developmental trajectories on each analyzed feature from the SPANGY decomposition. Fig. 2 depicts the normative trajectories for absolute band powers of frequency bands B4, B5, and B6.

**Fig. 2.**
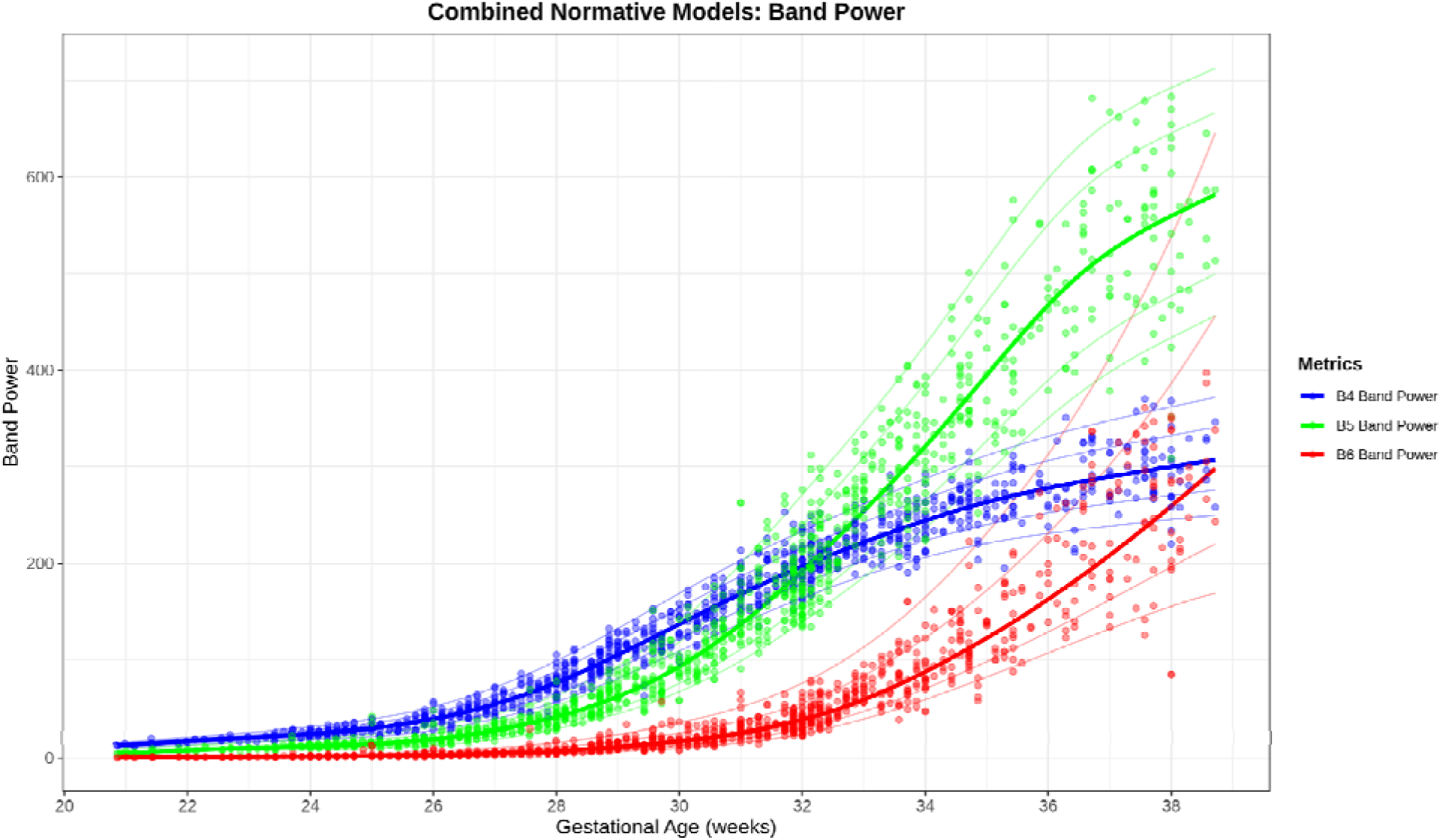
Normative curves for absolute band powers of B4 (blue), B5 (green), and B6 (red) across gestational ages from approximately 21 to 38 weeks, illustrating the typical developmental trajectories of spectral power for each frequency band. Individual data points from all subjects across the combined datasets are overlaid on the normative curves as colored dots, showing the distribution of observed values at each gestational age. The fitted curves represent the central tendency of each band’s developmental trajectory, while the lighter shaded regions surrounding each curve indicate the 3rd, 15th, 85th and 97th percentile intervals, capturing the expected range of normal variability.

The three bands demonstrate distinct developmental trajectories, differing in their onset timing, growth rates, and absolute magnitude of spectral power across gestation.

Band B4 (blue) shows the earliest emergence and follows a relatively linear developmental trajectory throughout the observed gestational period. The curve begins with detectable power values around 21-22 weeks, starting near zero and progressively increasing to approximately 300 units by 38 weeks. The data points show relatively tight clustering around the fitted curve at early ages (21 - 24 weeks), with band power values ranging from near zero to approximately 50 units at 26 weeks of gestational age . As gestational age advances, both the mean band power and the variability increase, with data points at 38 weeks of gestational age spanning from approximately 200 to 350 units. The prediction interval (light blue shaded region) widens progressively with advancing gestational age, reflecting increasing inter-individual variability in B4 power at older ages. The derivative (Shown on Supp. Fig. 3) confirms an inflection point at around 25 weeks, characterized by a steady increase up until 28 weeks where the growth of the curve begins to slow down.

Band B5 (green) demonstrates the steepest growth trajectory and reaches the highest absolute power values among the three bands at 38 weeks of gestational age. The curve shows minimal power values (near zero) at 21-24 weeks, followed by progressive acceleration that becomes particularly pronounced after approximately 30 weeks of gestational age. By 38 weeks, B5 band power reaches values of approximately 600 - 650 units, nearly double the maximum values achieved by B4 or B6. The individual data points show substantial scatter at mid-to-late gestational ages, with particularly wide dispersion visible from 32 weeks onward, where values at any given age can span a range of up to 100 units. The derivative (shown on Supp. Fig. 3) reveals inflection points around 25 and 29 weeks of gestational age, which is followed by an exponential increase in the growth rate up until 35 weeks, when it begins to slow down.

Frequency band B6 (red) demonstrates the most delayed onset, following an exponential growth trajectory comparable to B5, though with a reduced absolute magnitude. Power in B6 remains negligible until roughly 26-28 weeks, subsequently rising alongside an increasing growth rate that reaches its peak at approximately 38 weeks of gestational age (as illustrated on Supp. Fig. 3). The growth rate also reveals an inflection point at 30 weeks of gestational age. By term age (38 weeks), B6 reaches values of approximately 300 units, similar in magnitude to B4 despite its later emergence. The data points for B6 show the tightest clustering at early gestational ages and progressively increasing scatter as gestation advances, with the prediction interval (light red shaded region) widening substantially after 32 weeks of gestational age. Across all three bands, the normative curves successfully capture the central tendency of the observed data points, with the majority of individual observations falling within the prediction intervals at all gestational ages.

Unlike the absolute band power trajectories, the surface area percentage curves (Fig. 3) reveal fundamentally different developmental patterns, with B4 displaying an inverted U-shaped trajectory, while B5 and B6 show increasing trends with distinct onset timings and growth rates.

**Fig. 3.**
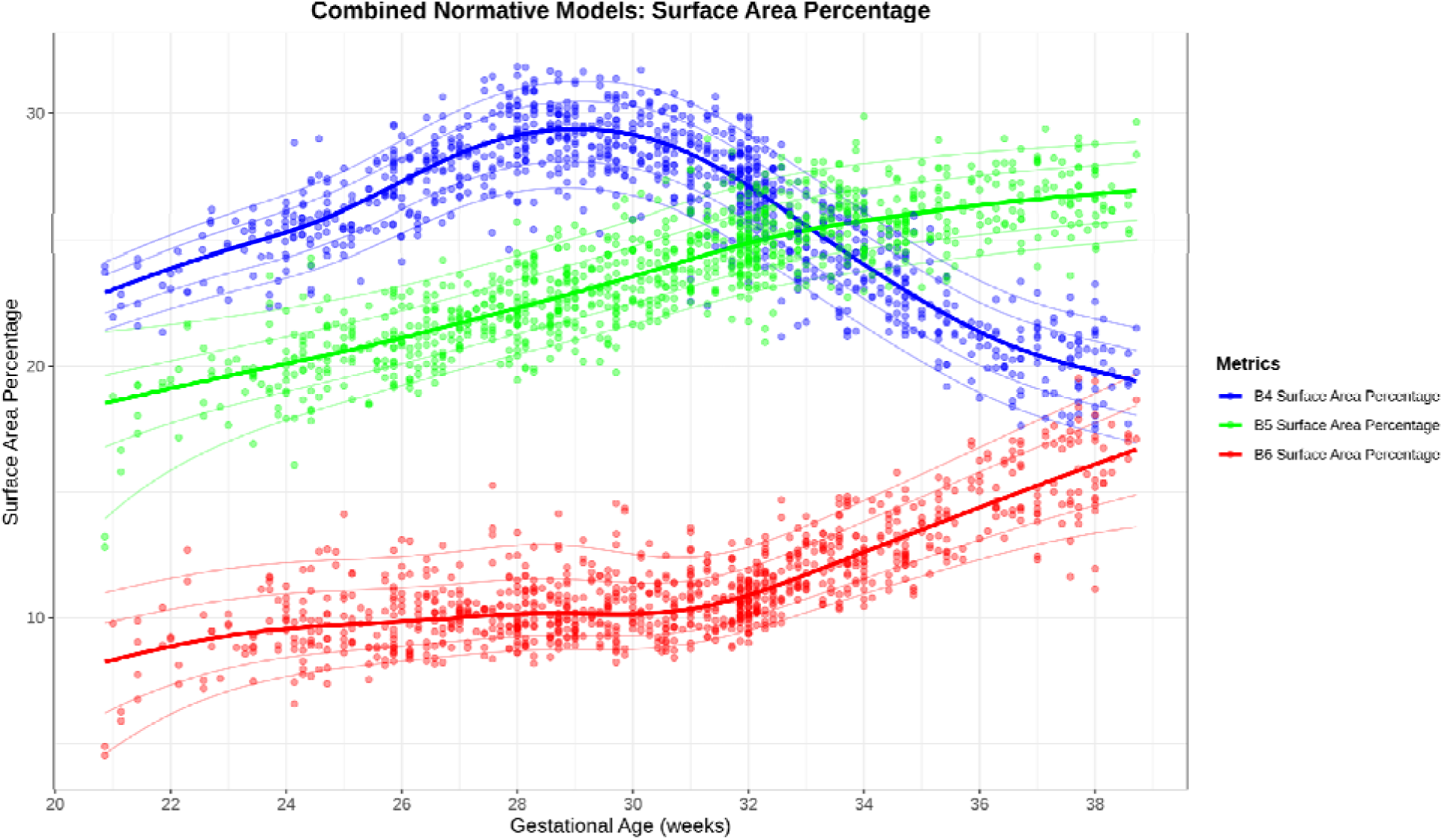
Normative curves for the percentage of cortical surface area dominated by bands B4 (blue), B5 (green), and B6 (red) across gestational age from approximately 21 to 38 weeks. Individual data points from all subjects across the combined datasets are overlaid as colored dots, showing the distribution of observed percentage values at each gestational age. The fitted curves represent the central tendency of each band’s surface area contribution, while lighter shaded regions surrounding each curve indicate the 3rd, 15th, 85th and 97th percentile intervals.

Band B4 (blue) exhibits a distinctive biphasic trajectory characterized by an initial increase followed by a subsequent decline. At 20-21 weeks of gestational age, B4 accounts for approximately 22-23% of the cortical surface area. The percentage increases progressively through early-to-mid gestation, reaching a peak of approximately 28-29% around 28-30 weeks. Following this peak, B4 surface area percentage begins to decline steadily, decreasing to approximately 26-27% by 32 weeks, 23-24% by 35 weeks, and reaching approximately 18-19% by 38 weeks. The individual data points show considerable scatter throughout the gestational range, with particularly wide dispersion visible at the peak period (27-31 weeks) where values span from approximately 25% to 33%. The prediction interval (light blue shaded region) maintains relatively consistent width throughout the trajectory, widening slightly during the peak period and the later decline phase.

Band B5 (green) demonstrates an increasing trajectory throughout gestation, starting from approximately 14-15% surface area coverage at 21 weeks of gestational age and increasing progressively to approximately 26-27% by 38 weeks. The growth pattern appears relatively linear through early-to-mid gestation (20-32 weeks), with percentage values increasing from around 15% to 23-24%. After 32 weeks, the rate of increase appears to decelerate slightly, with the curve approaching a value around 26-28% by term age. Individual data points show substantial variability at all gestational ages, with scatter increasing noticeably after 28 weeks where values at any given age can span a range of 8-10 percentage points. The prediction interval (light green shaded region) widens progressively with advancing GA, particularly after 32 weeks of gestational age.

Band B6 (red) exhibits a latest onset and follows a different growth pattern. At 21-22 weeks of gestational age, B6 accounts for only 3-5% of cortical surface area, with values remaining relatively flat below 8-9% through approximately 26-27 weeks of gestational age. After this initial plateau phase, B6 percentage begins to increase with accelerating growth rate, rising to approximately 10% by 30 weeks of gestational age, 12-13% by 34 weeks of gestational age, and reaching approximately 17% by 38 weeks of gestational age. The individual data points for B6 show the tightest clustering at early gestational ages (20-26 weeks of gestational age) where values are uniformly low, with increasing scatter visible after 28 weeks of gestational age as the growth phase begins. The prediction interval (light red shaded region) remains narrow through the early plateau phase and widens considerably during the accelerated growth period after 30 weeks of gestational age, reflecting increasing inter-individual variability as B6 contributions expand.

Fig. 4 shows the relative band powers across bands B4-B6. Unlike absolute band power, which measures spectral magnitude in arbitrary units, relative power values are normalized proportions ranging from 0 to 1, reflecting how the total folding energy is distributed across different spatial scales at each developmental stage and serve as complimentary metrics to the surface area percentage. The overall trend of the three bands are similar to those previously described under the surface area percentage.

**Fig. 4.**
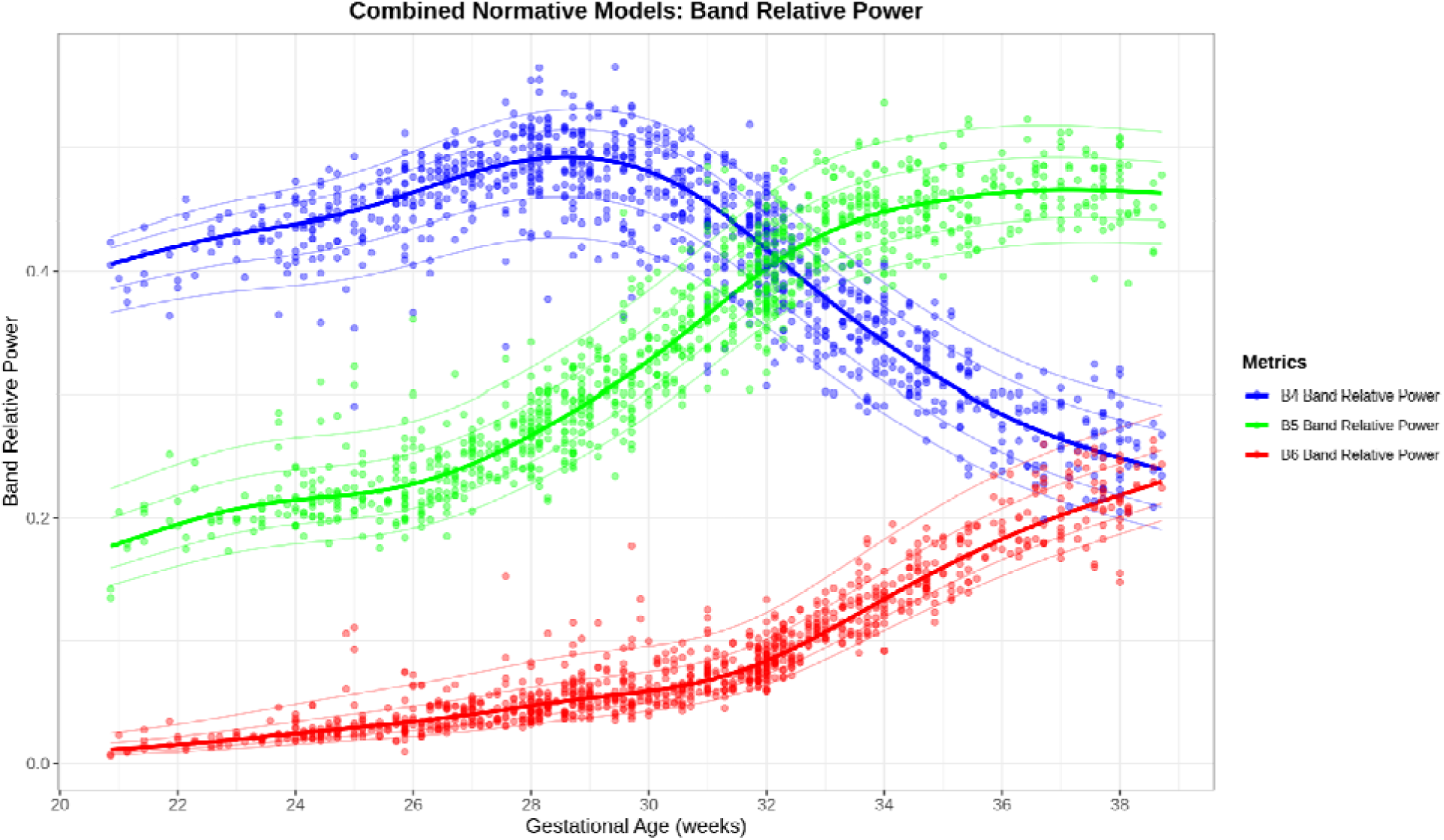
Normative curves for relative band powers of B4 (blue), B5 (green), and B6 (red) across gestational age from approximately 21 to 38 weeks of gestational age, where relative power represents each band’s proportional contribution to the total folding power across all frequency bands. Individual data points from all subjects across the combined datasets are overlaid as colored dots, with fitted curves representing the central tendency and lighter shaded regions indicating the 3rd, 15th, 85th and 97th percentile intervals.

### 3.2. Frequency bands , qualitative, visualization

The result of the spectral decomposition of the cortical surface curvature was used to create a dominant band segmentation map. Each vertex on the cortical mesh was assigned to its dominant frequency band based on the spectral power contribution at that location, resulting in a color-coded visualization as seen on Fig. 5.

**Fig. 5.**
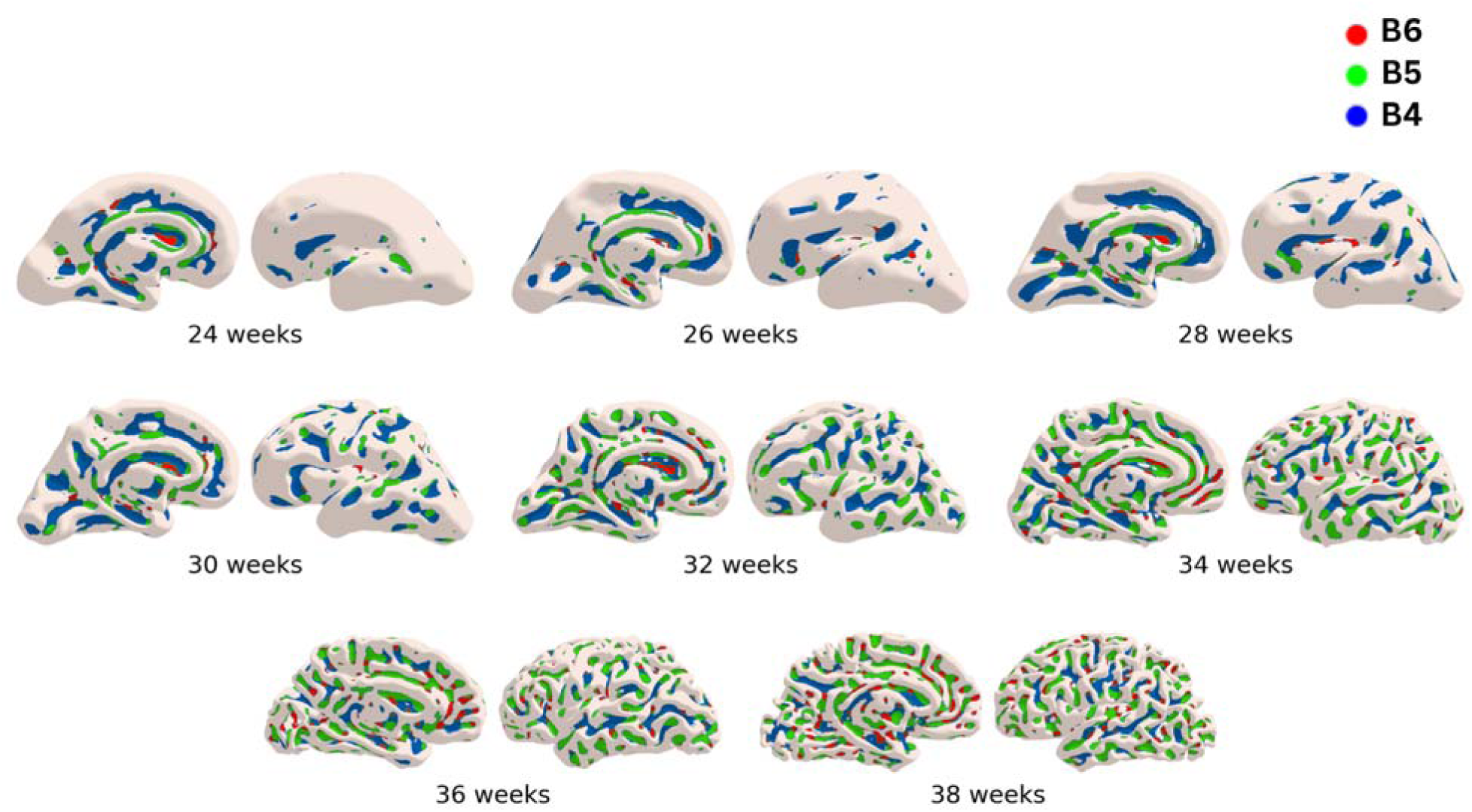
displays the dominant frequency band segmentation map and the developmental progression of frequency bands B4, B5, and B6 across the entire cortical hemisphere from 24 to 38 weeks gestational age. Each vertex on the cortical mesh was assigned to its dominant frequency band based on the spectral power contribution at that location, resulting in a color-coded visualization that illustrates the partitioning of the cortical surface into distinct folding domains according to spatial scale (blue for B4, green for B5, red for B6). All images have been scaled independently to emphasize developmental transitions rather than to represent uniform dimensions.

The visualization of the dominant band at every vertex at term (Fig. 5) reveals several notable spatial patterns in the distribution of frequency bands. Band B4 (blue) occupies large, continuous regions of the cortical surface, particularly visible in the depths of major sulci including the superior temporal sulcus, central sulcus, and other primary anatomical landmarks. These B4-dominant regions form broad swaths across the surface of the hemisphere, with the blue patches extending deeply into sulcal valleys and across wide gyral expanses. The spatial extent of B4 regions are mostly located in the large-scale folding patterns of the cortex.

Band B5 (green) appears in more spatially restricted regions compared to B4, forming intermediate-sized patches distributed across the cortical surface. Green patches are particularly prominent along the walls and margins of sulci, often forming transition zones between the deep B4-dominant sulcal floors and the superficial cortical surface. The distribution of B5 shows a more heterogeneous spatial pattern than B4, with multiple scattered green patches rather than large continuous territories.

Band B6 (red) is present as small, discrete patches scattered across the cortical surface, representing the finest spatial scale of folding captured by the SPANGY decomposition. Red patches appear as localized spots, often situated at the edges of sulci, on narrow gyral banks, or in areas of complex, high-curvature geometry. The B6 regions are notably less extensive than either B4 or B5, appearing as punctate features rather than large contiguous areas. These small red patches are distributed throughout the cortex but show particular concentration in regions with elaborate surface convolutions.

In order to understand the formation of such patterns throughout fetal development, Fig. 6 shows the spatial distribution of dominant bands within a specific region (the central sulcus) between 26w and 38w of gestational age, after smoothing the cortical surface for better visualization.

**Fig. 6.**
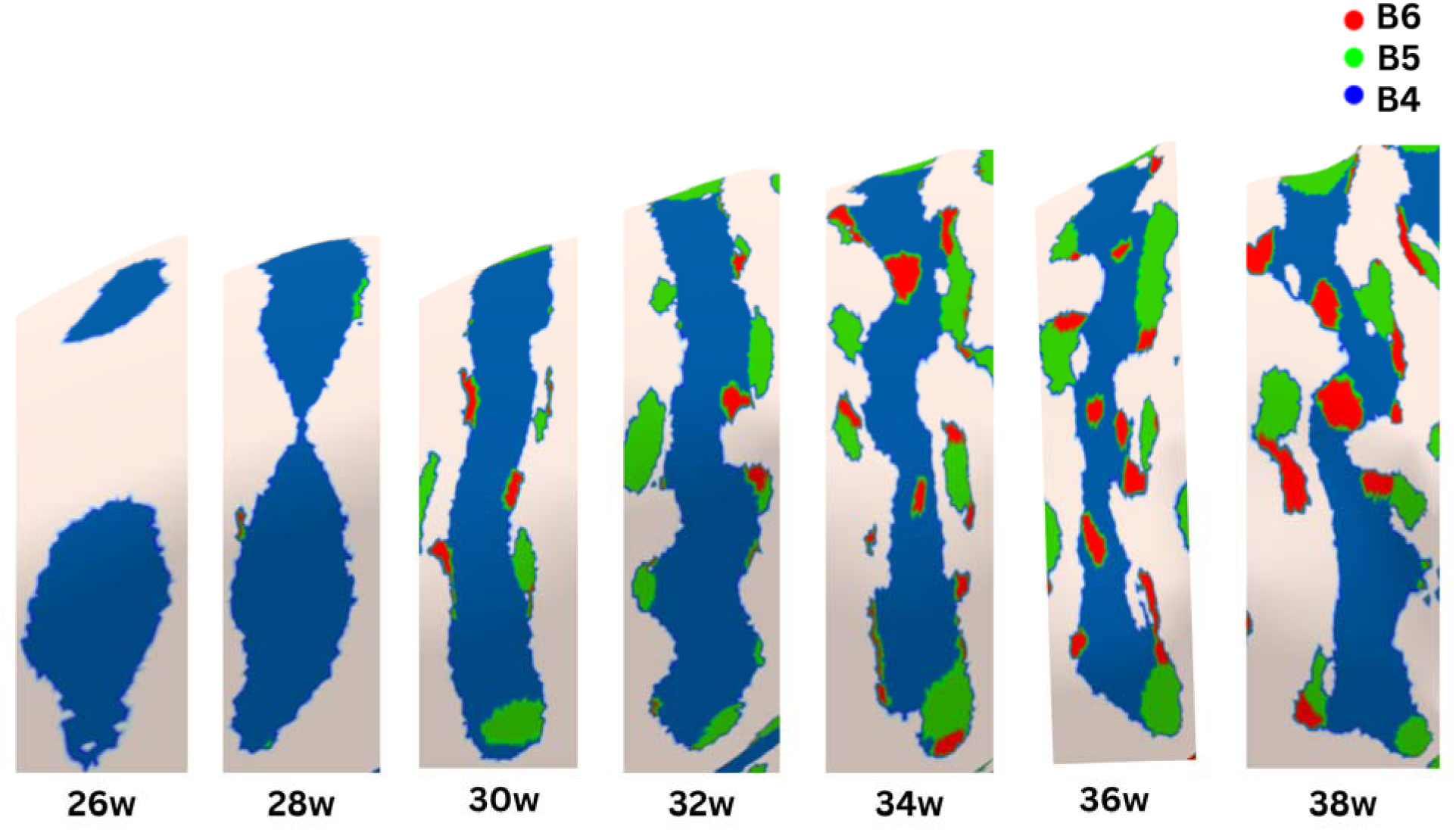
presents a developmental time series showing the spatial distribution of dominant frequency bands (B4, B5, and B6) within the central sulcus region across seven gestational ages from 26 to 38 weeks. Each panel displays a close-up view of the same anatomical region, with vertices color-coded according to their dominant spectral band: blue for B4, green for B5, and red for B6. Note that the panels have been enlarged to highlight the developmental changes and are not displayed at a uniform scale.

On Fig. 6, we observe that at 26 weeks of gestational age, the central sulcus region displays almost exclusive B4 dominance, with blue patches occupying the entire visible sulcal area as a single continuous domain. By 28 weeks, the sulcal region remains predominantly B4-dominant, though small green (B5) patches begin to appear at the superior margin of the sulcal bank, marking the first emergence of intermediate-scale folding contributions.

From 30 to 34 weeks of gestational age, progressive diversification of frequency band composition becomes evident. At 30 weeks, B5 patches appear along both sulcal margins as discrete clusters, while small red (B6) spots make their first appearance near high-curvature regions. The 32-week panel shows B5 occupying substantially larger regions along the sulcal banks as both continuous strips and discrete clusters, while B6 patches increase in number and area. By 34 weeks, green (B5) territories have expanded to cover substantial portions of the sulcal banks, appearing as large irregularly shaped regions interspersed with blue B4 domains, while red (B6) patches are distributed throughout the visible area, particularly at interfaces between B4 and B5 regions.

At 36 and 38 weeks of gestational age, the spatial distribution of the frequency bands demonstrates high complexity. The 36-week panel shows a complex mosaic pattern where B4 no longer forms a continuous dominant territory but persists in the deep sulcal floor and larger patches, while B5 occupies multiple large patches across sulcal banks and B6 appears as numerous small discrete regions concentrated in geometrically complex zones. By 38 weeks, the three bands form an intricate interwoven pattern, where B4 persists in discrete patches primarily along the sulcal floor, B5 occupies extensive irregular territories across the sulcal banks, and B6 appears as numerous small spots concentrated at high-curvature locations. Across the entire 26-38 week sequence, the initial exclusive B4 dominance progressively transitions to increasing B5 and B6 contributions, with B5 first appearing at margins and expanding inward, while B6 emerges later and concentrates in areas of highest geometric complexity.

## 4. Discussion

This study represents the first application of Spectral Analysis of Gyrification (SPANGY) to multi-centric fetal brain imaging data. By extending SPANGY methodology, which was originally developed and validated in adult and pediatric populations, to the fetal developmental window, we have explored a possibility for quantitative characterization of early cortical development. We release the first normative curves of frequency decomposition of cortical folding development during the prenatal period that could serve as normative references in future application studies.

Our dataset constitutes the largest fetal brain cohort analyzed to date, encompassing 635 subjects across a gestational age range of 20–38 weeks from multiple imaging sites. The multi-center nature of this dataset inherently provides greater generalizability and robustness compared to single-center investigations, as it captures natural variability across different populations, scanning protocols, and scanner manufacturers. This diversity enhances the external validity of our normative models and increases confidence that the developmental trajectories we observe reflect true biological patterns rather than site-specific artifacts or sampling biases.

### Normative Modeling and Harmonization

Our analysis of site effects revealed significant variance in raw SPANGY metrics, necessitating statistical harmonization using ComBat-GAM (Pomponio et al., 2020) before model selection. This harmonization approach removes site-related technical variance while preserving biological variability associated with gestational age and individual differences, ensuring that our normative curves reflect true developmental patterns rather than varying differences from acquisition parameters (Bethlehem et al., 2022).

The choice of GAMLSS framework with SHASH distribution was motivated by the non-Gaussian, heteroskedastic nature of neuroimaging data acquired during early brain development. Fetal brain metrics often exhibit skewness and kurtosis that violate assumptions of simpler modeling approaches, and variance typically changes across gestational age as inter-individual variability increases with progressive cortical differentiation. Our systematic model comparison procedure, progressing from linear models through GAMs to full GAMLSS, ensured that we selected models with appropriate complexity to capture developmental trajectories without overfitting. The model selection criterion using the log score of the test set, in a cross-validation setting, provided protection against both underfitting (insufficient model complexity) and overfitting (excessive complexity that degrades generalization) ensuring that the models exhibited goodness of fit on the data.

The generated normative trajectories effectively characterize the central tendency of the observed data, demonstrating concordance with expected developmental patterns. As depicted in the band power trajectories (Fig. 2), band B4 exhibits the most constrained variability among the three primary folding bands (B4, B5, and B6). This observation is consistent with the expectation that inter-individual variability in cortical morphology increases with folding complexity (Akula et al., 2023). This phenomenon is accurately represented within our normative framework, particularly for bands B5 and B6, which demonstrate expanded prediction intervals at advanced gestational ages, reflecting the heightened geometric heterogeneity inherent to more complex folding patterns.

Critically, the normative models reveal not only when each fold scale emerges but also how their relative contributions evolve throughout pregnancy. These could serve as valid comparisons for evaluating novel analysis techniques and for validating computational models of cortical morphogenesis. Furthermore, the scale-specific growth patterns we have quantified could provide empirical constraints for biomechanical models attempting to simulate the folding process, offering testable predictions about development at different spatial scales.

### Situating our new results with respect to previous SPANGY studies

Our findings demonstrate notable consistency with previous SPANGY-based investigations of cortical folding development. Specifically, our fetal-period trajectories establish developmental continuity with the postnatal trajectories reported in Dubois et al., 2019, as evidenced by comparable patterns in the normative curves of the relative band power proportion dynamics. For both studies, at 38 weeks of gestational age (38w post-menstrual age in Dubois et al., 2019), the relative band powers observed were approximately 0.2 for frequency band B4, 0.45 for B5, and 0.2 for B6. This suggests a conserved mechanism of cortical folding across the perinatal transition.

However, systematic differences in absolute band power magnitudes were observed between our fetal cohort and the postnatal cohort of (Dubois et al., 2019), with higher values reported in the latter. This discrepancy may reflect either inter-site measurement variability, or genuine neurobiological effects associated with the perinatal transition, including the documented impact of birth on gyrification (Mihailov et al., 2025).

The normative trajectories established in this study provide a quantitative reference framework for investigating aberrant cortical folding in pathological populations. Such comparisons have proven valuable in prior work, as demonstrated in (Germanaud et al., 2014) in the context of severe developmental microcephalies, and our prenatal reference data now enable an extension of such investigations to earlier developmental windows when cortical malformations may first emerge.

### Dynamics of Frequency Bands in Fetal Development

Our analysis of frequency bands B4, B5, and B6 in fetal brains provides some insight into the hierarchical organization of early cortical folding. Band B4 captures the earliest-emerging and most prominent sulcal structures, including major sulci such as the central sulcus, and superior temporal sulcus. These primary structures form the fundamental organizational framework of the cortical surface and exhibit robust presence even in younger gestational ages within our sample. B4 frequency band power increases steadily across gestational age, reflecting the progressive deepening and elaboration of primary sulci as the cortex matures.

Band B5 represents folding patterns at intermediate spatial scales. This also includes sulci that emerge later in development, falling between the major primary sulci and the finer tertiary patterns. B5 frequency band power shows a similar upward trajectory as that of B4 but with a temporal lag, consistent with secondary folds emerging after primary structures have been established. Band B6 captures the finest folding details, which represent the last folding patterns to emerge and show the greatest inter-individual variability. B6 frequency band power exhibits the most dramatic increases in the later gestational ages, corresponding to the emergence and proliferation of tertiary folding patterns in the third trimester.

### Surface Area and Relative Band Power Interpretations

The proportional surface area occupied by each frequency band offers complementary insights into the spatial distribution of emerging folding patterns. As illustrated in Fig. 3, the expansion of the B4 surface area coverage corresponds to the morphological establishment, growth, and progressive deepening of primary sulci and fundamental neuroanatomical landmarks. Since these structures constitute the structural foundation for subsequent gyrification, they exert near-complete dominance over the cortical surface area during early gestational periods. The initial deceleration in the expansion rate of B4 coverage occurs in temporal alignment with the accelerated growth of the B5 surface area, signifying not only the sustained volumetric expansion and geometric complexification of the cortex but also a systematic redistribution of territories previously characterized by B4 dominance. Consequently, band B5 represents both the emergence of novel sulcal structures and the localized geometric elaboration of existing primary sulcal regions. A subsequent, more pronounced decline in the proportional surface area of B4 is observed to coincide with the rapid expansion of B6 coverage, further substantiating the redistribution of both B4 and B5 territories driven by the escalating hierarchical complexity of cortical folding. This shift reflects the increasing sophistication of the cortical landscape as higher-order folding levels are superimposed upon the primary structural scaffold. Fig. 6 illustrates the emergence of B5 and B6 band patches specifically on sulcal walls in regions already committed to folding at lower frequencies. This further demonstrates that the folding complexity continues to increase locally even after initial fold formation, generating increasing geometric complexity that suggests that there are different folding types within a single fold. This further supports the idea that gyrification happens in successive waves of folding (Dubois et al., 2019). The systematic temporal progression observed, with B4 emerging earliest and dominating early gestation, followed by a progressive expansion of B5 and later emergence of B6, allows us to access a finer-scale description of folding at the level of sulci than the established sequence of primary, secondary, and tertiary fold formation documented in classical anatomical studies (Chi et al., 1977; De Vareilles et al., 2023; Feess-Higgins and Larroche, 1987).

This developmental dynamic is linked to gestational age, brain size, and the mechanical constraints of brain tissue growth as inferred by biomechanical models (Bayly et al., 2014; Tallinen et al., 2016), substantiating that differential volumetric expansion of brain tissues generates increasingly complex folding patterns at progressively finer spatial scales, and corroborating the claim that larger cortical surfaces exhibit inherently greater gyrification complexity (Germanaud et al., 2012).

Relative band power metrics characterize the proportional contribution of each hierarchical folding level to the global cortical architecture, providing quantitative insights into the dynamic interplay between spectral frequency components across the developmental window. Observations derived from our analysis indicate that the relative proportions of frequency components undergo substantial reconfiguration throughout the prenatal period of cortical maturation. Fig. 4 illustrates the recurring developmental trend where early gestational ages are characterized by a marked dominance of B4 power, which is progressively superseded by increasing contributions from bands B5 and B6 as gestation advances. These findings effectively complement the surface area percentage results, suggesting a robust coupling between the spatial extent and the spectral contribution of specific folding scales. Furthermore, these metrics facilitate reliable cross-study comparisons even in the absence of explicit harmonization, as the proportional distribution of folding energy across spatial scales is largely invariant to site-specific acquisition parameters. This methodological consistency is exemplified by the high degree of concordance observed between our fetal results and the postnatal values reported in (Dubois et al., 2019).

## Clinical Application

The fine-grained quantification of gyrification dynamics offered by SPANGY, and the normative curves established in this study, hold some promise for earlier detection of subtle anomalies in cortical folding patterns. These data could facilitate the identification of deviations in gyrification that may escape detection using current imaging techniques, potentially enabling the validation of early biomarkers for neurodevelopmental disorders of genetic origin. For instance, genetic syndromes associated with subtle but pathognomonic gyrification abnormalities, such as Williams syndrome, could be flagged earlier in pregnancy, allowing for timely genetic counseling and intervention planning (Kippenhan et al., 2005; Thompson et al., 2005). This approach may be particularly valuable for conditions where early brain development is affected by genetic mutations that disrupt cellular processes (e.g., neural migration or proliferation), which in turn alter the mechanical forces driving cortical folding.

Furthermore, SPANGY’s multi-scale analysis could prove invaluable for studying the impact of prenatal environmental factors on cortical development. Situations such as intrauterine growth restriction (IUGR) (Dubois et al., 2008; Egaña-Ugrinovic et al., 2013), maternal stress (Wu et al., 2020), or exposure to teratogens, congenital heart disease (Brossard-Racine et al., 2016; Juergensen et al., 2024), may leave distinct signatures in the spectral profiles of gyrification, reflecting how adverse conditions perturb the hierarchical emergence of cortical folds. By comparing SPANGY-derived metrics in fetuses exposed to such conditions against normative trajectories, researchers could elucidate how specific prenatal insults selectively affect different scales of cortical folding, whether by delaying the onset of primary sulci, altering the proportional contribution of intermediate (B5) or fine-scale (B6) patterns, or disrupting the temporal coordination between folding scales. This could, in turn, provide mechanistic insights into the pathways linking prenatal adversity to long-term neurodevelopmental outcomes.

## Limitations and Future Directions

Several limitations warrant consideration. Firstly, despite the advantages of multi-center datasets, the integration of data from different scanners and acquisition protocols introduces substantial challenges. Scanner-specific differences in spatial resolution, signal-to-noise ratio, contrast characteristics, and geometric distortion can systematically bias morphometric measurements and confound apparent site effects with true biological differences. While our harmonization procedures address much of this variance, it is impossible to fully disentangle technical from biological sources of variation in observational multi-site studies. Residual site-related variance may persist even after harmonization, potentially introducing subtle biases in our normative curves or increasing uncertainty in model predictions. Though we believe that this is not the case in our study.

Secondly, our cross-sectional design precludes direct observation of individual developmental trajectories; longitudinal studies would provide more definitive characterization of within-subject folding dynamics. Furthermore, while our dataset is large by fetal neuroimaging standards, the sample size at specific gestational ages may limit statistical power for detecting subtle developmental inflections or nonlinearities.

Thirdly, our normative models assume that our sample represents typical development, but systematic biases in recruitment or inclusion criteria could affect generalizability, although the multicentric aspect of our study and the general agreement across centers suggest little or no such bias.

Lastly, small-scale noise, reconstruction artifacts, or segmentation errors can introduce random high-frequency components that may be misattributed to genuine folding patterns. This suggests that our higher-frequency band measurements (particularly B6) may contain higher noise-related variance than lower-frequency bands (B4 and B5). To address these technical concerns, this study employed a rigorous quality control protocol intended to minimize the influence of mesh irregularities on the final analysis.

Future investigations should incorporate automated mesh quality assessment and spectral methods that are more robust to noise. This could further improve the reliability of SPANGY metrics in challenging fetal imaging scenarios. Future studies should also extend the analysis to include clinical populations to assess sensitivity to neurodevelopmental pathology, and explore associations between SPANGY metrics and functional or cognitive outcomes in longitudinal follow-up studies such as (Ji et al., 2024).

## 5. Conclusion

In conclusion, this work represents a significant advance in quantitative prenatal neuroimaging, demonstrating that spectral analysis can successfully characterize the complex, multi-scale process of fetal cortical folding and provide validated normative references that enable statistically rigorous assessment of individual developmental trajectories. By bridging the gap between qualitative anatomical descriptions and quantitative developmental neuroscience, this framework opens new avenues for investigating the origins of cortical architecture and for detecting early signatures of atypical neurodevelopment.

## 6. Data Availability

Data for fetal subjects used in the preparation of this manuscript, from the developing Human Connectome Project (dHCP) cohort, is publicly available (https://biomedia.github.io/dHCP-release-notes/). We also used fetal data from 3 other sites (MarFet, BCNatal and CHUV); however, to preserve and protect the identities of the subjects, their data is not currently openly available. It can, however, be accessible upon reasonable request.

## 7. Code Availability

All relevant codes relating to the methods used in our manuscript are publicly available at fetpype: https://github.com/fetpype/fetpype, and SLAM: https://github.com/brain-slam/slam.

## 9. Acknowledgments

## Ethical Approval

Ethical approval was obtained from all centers (La Timone: Aix-Marseille University N°2022-04-14-003; BCNatal: HCB/2022/0533; CHUV: CER-VD 2021-00124)

## 10. Author Contributions

Framing research questions, study design and writing the first version of the paper: Harvey Dienye, Angeline Mihailov, Nadine Girard, Julien Lefèvre, Olivier Coulon, and Guillaume Auzias.

Data acquisition: Léo Pomar, Joanna Sichitiu, Vincent Dunet, Mériam Koob, Elisenda Eixarch, Aurelie Manchon, Nadine Girard, Mathieu Milh

Data processing and data quality check, : Harvey Dienye, Thomas Sanchez, Gerard Martí-Juan, Raquel G López.

Analytical and contextual contributions: all authors Manuscript editing: all authors.

Final approval of manuscript: all authors.

## Supplementary Materials

**Supp. Fig. 1.**
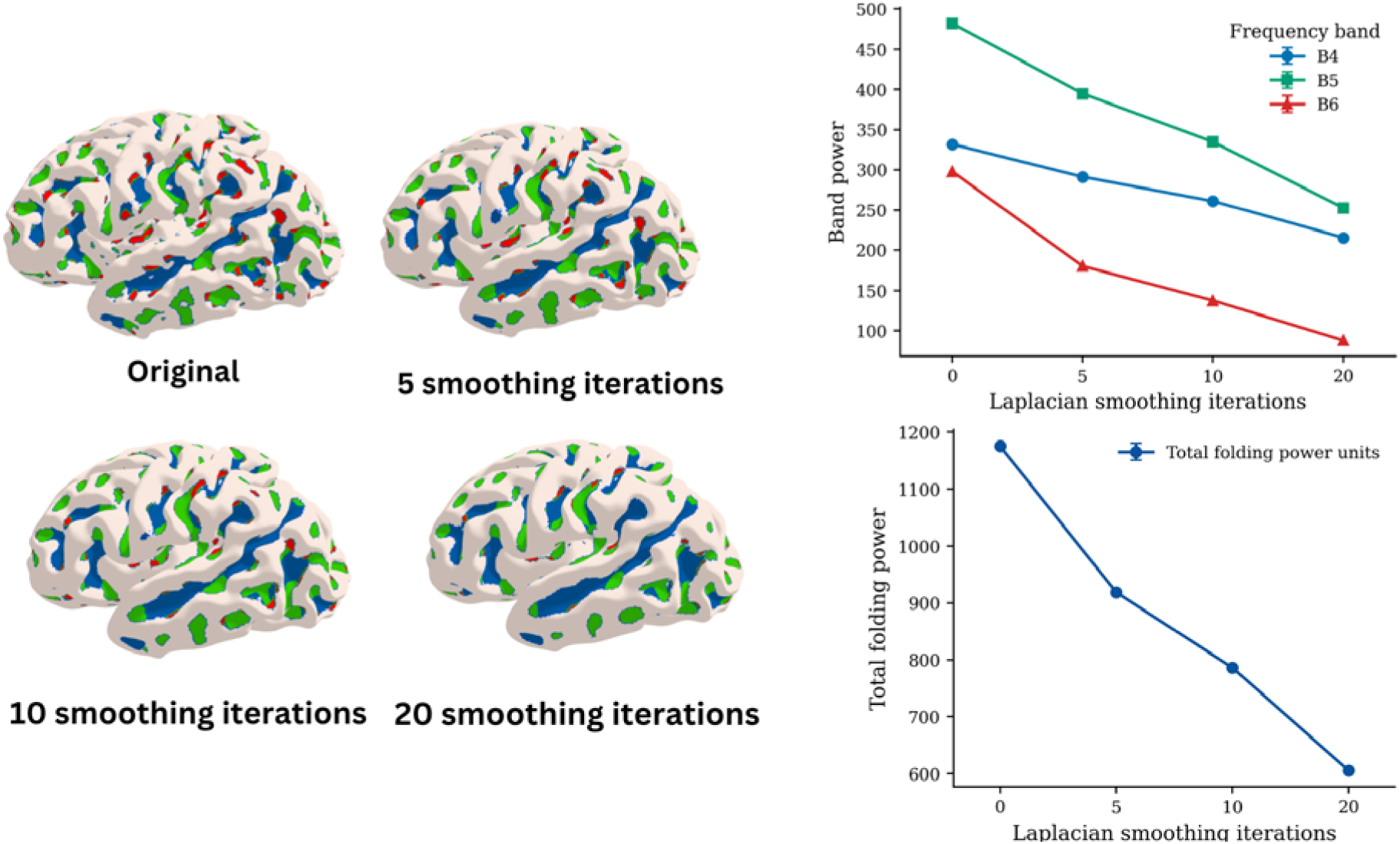
Relationship between the number of smoothing iterations, absolute band powers for B4-B6, and the total folding power on a representative fetal subject.

**Supp. Fig. 2.**
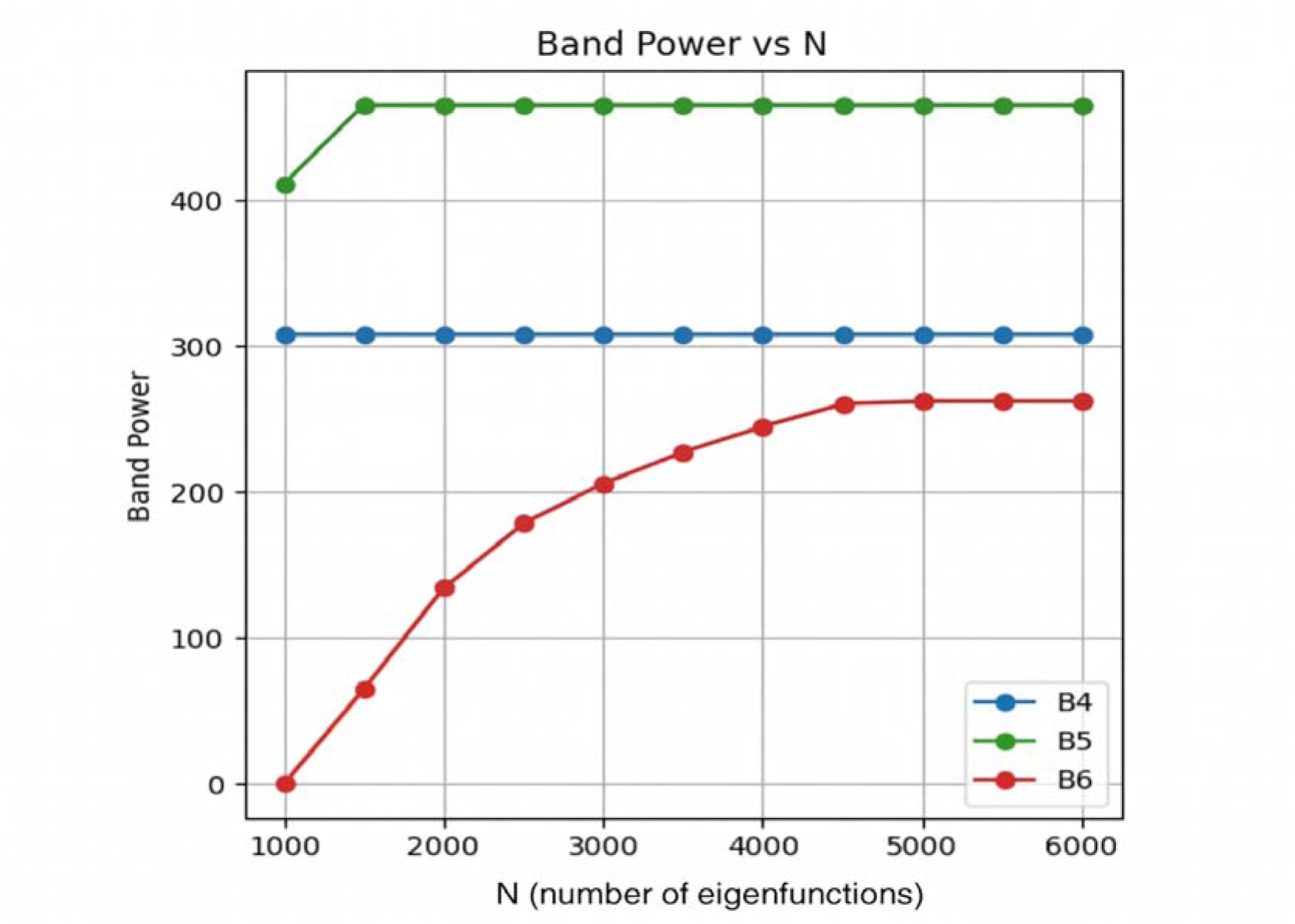
Relationship between the number of eigenfunctions (N) and the absolute band powers for B4-B6 on a representative fetal subject.

**Supp. Fig. 3.**
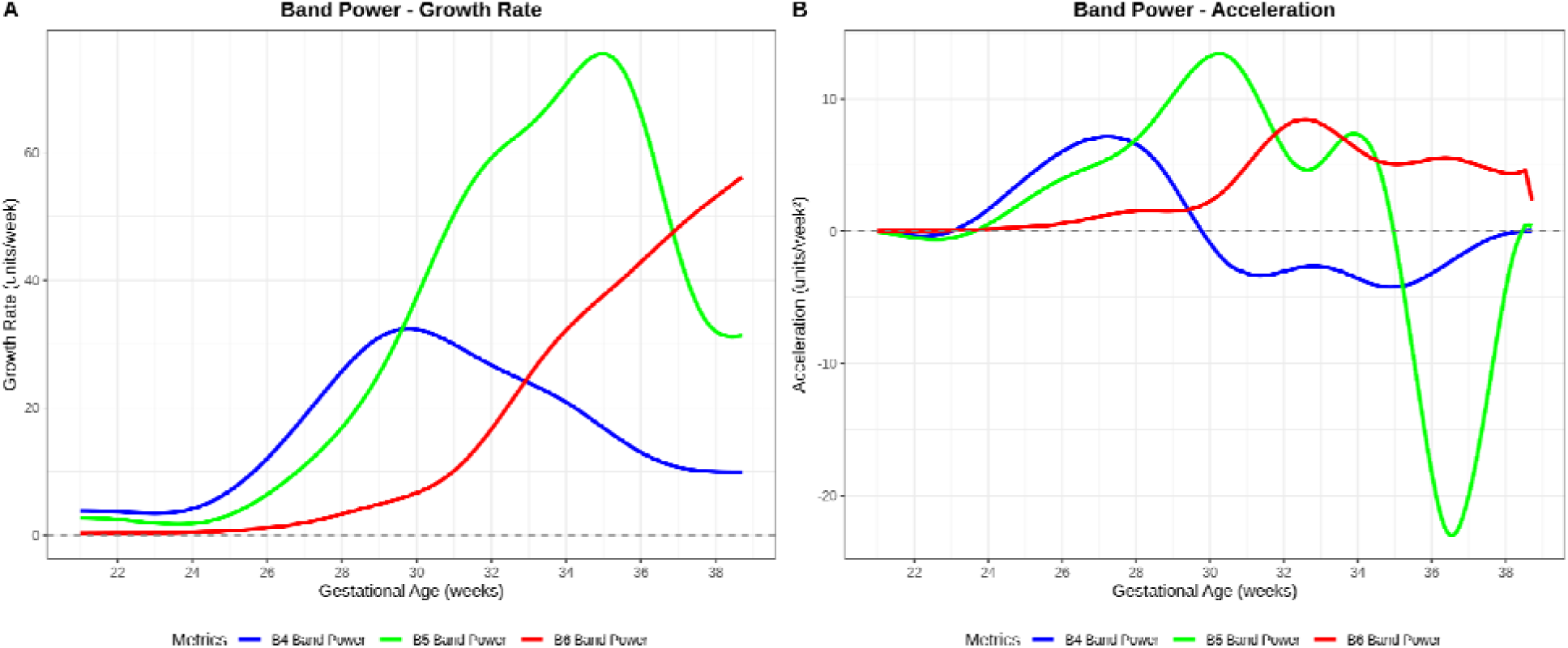
Growth rate (first order derivative) and acceleration (second order derivative) of the absolute band powers B4 - B6 across all gestational ages.

**Supp. Fig. 4.**
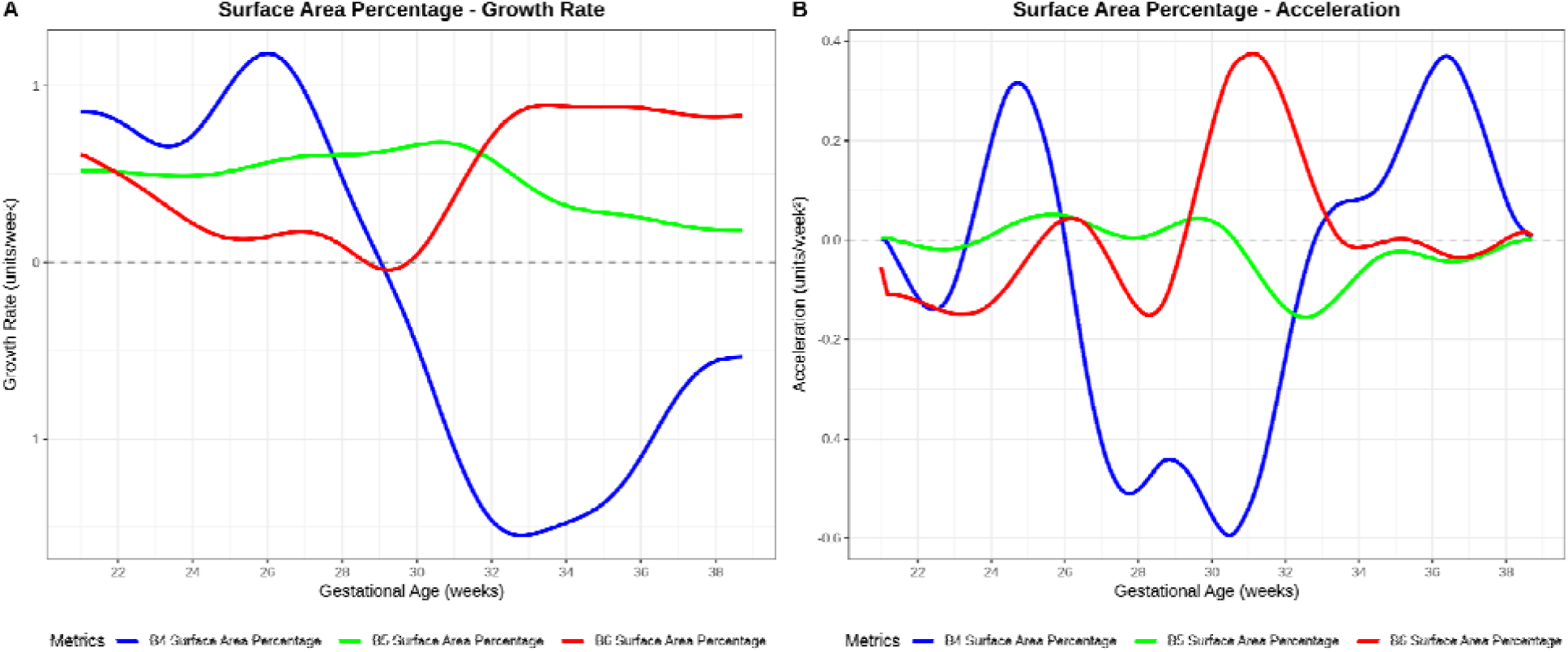
Growth rate (first order derivative) and acceleration (second order derivative) of the percentage surface area of B4 - B6 across all gestational ages.

**Supp. Fig. 5.**
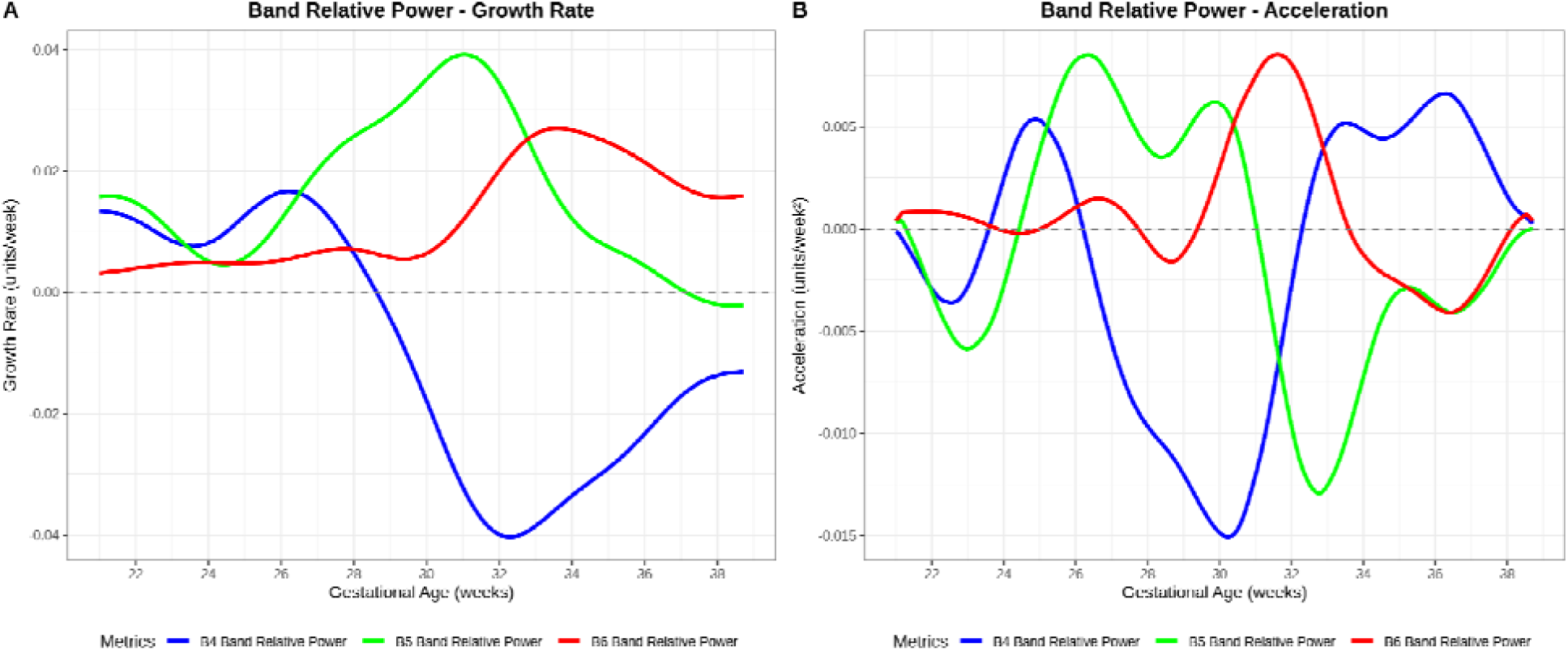
Growth rate (first order derivative) and acceleration (second order derivative) of the relative band powers B4 - B6 across all gestational ages.

**Supp. Fig. 6.**
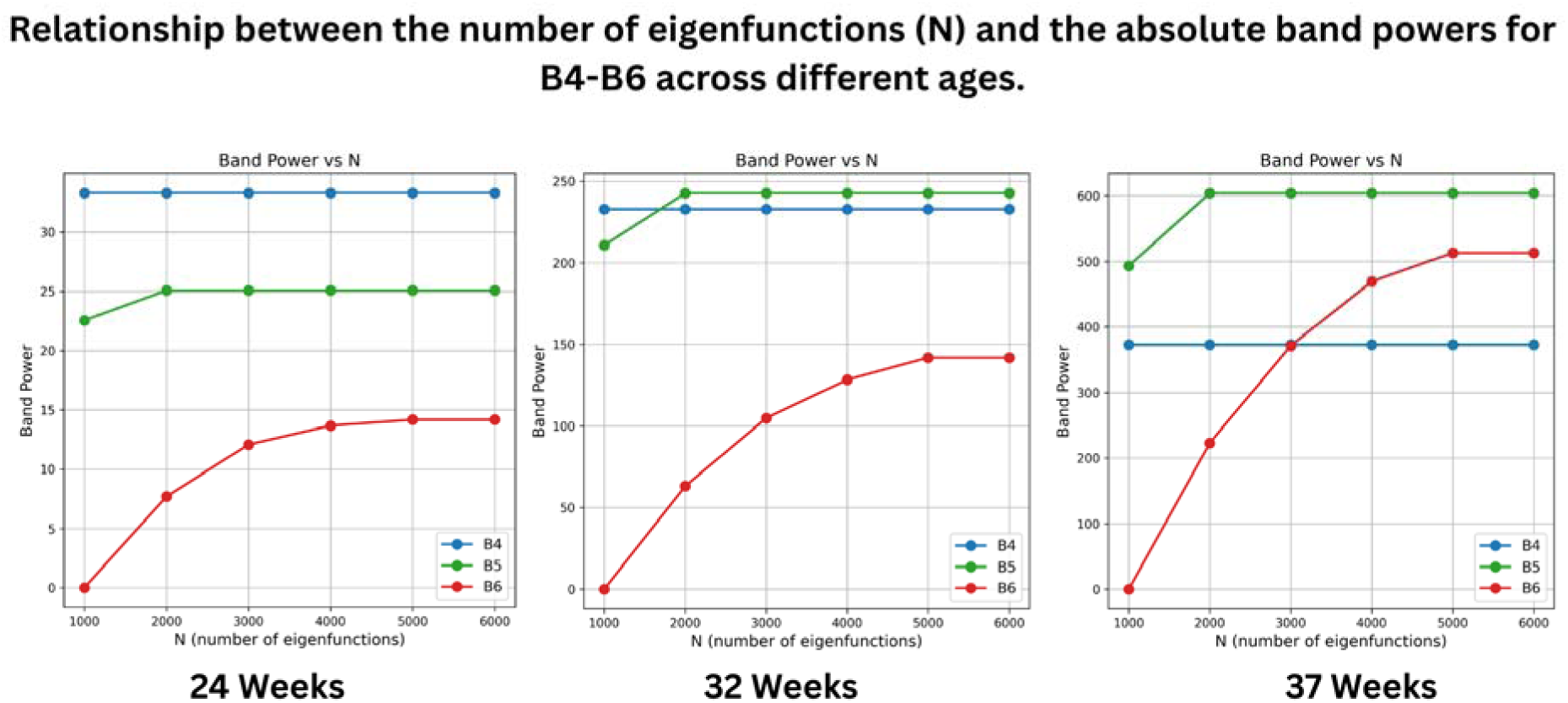
The eigenfunction calculation is influenced by the number of vertices. The higher the number of vertices, the more eigenfunctions can be calculated (up until the max number of vertices). With this in mind, we ran the computation on a mesh whose number of vertices is in the 95th percentile of the dataset. A plateau of band power B6 was seen at 5000. This means that band power B6 on meshes whose number of vertices are less than the chosen mesh would also plateau at 5000 or below.

**Supp. Fig. 7.**
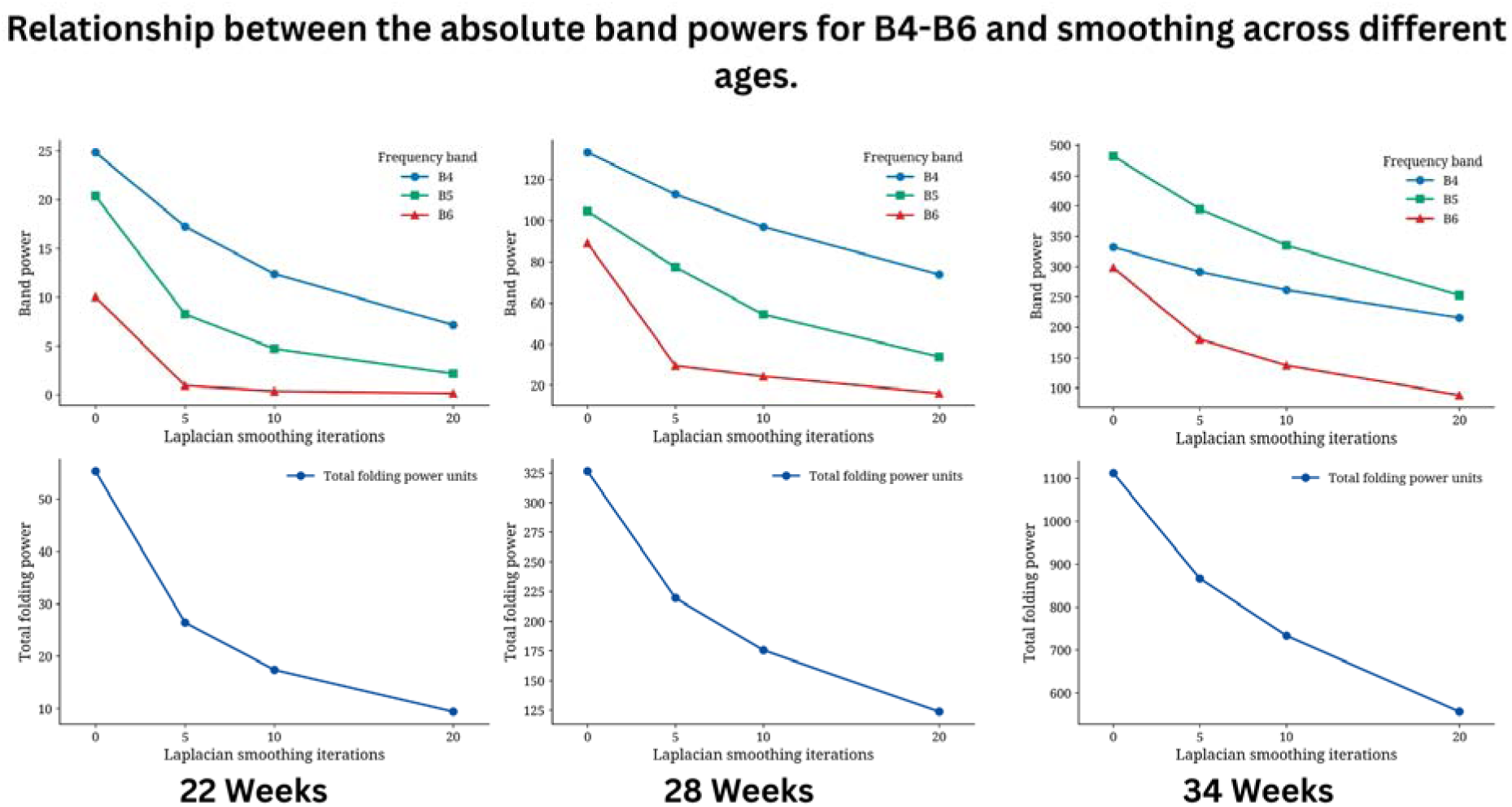
A noticeable elbow is observed after 5 smoothing iterations, across ages. It was also visually confirmed on Supp. Fig. 1. in the manuscript, to be the point after which we begin to lose anatomic features.

**Supp. Table 1.** Results of the analysis of the site effect performed for each feature extracted after SPANGY analysis.

| Feature | p_value |
| --- | --- |
| **B4 Surface Area Percentage** | 5.22066657765205e-11 |
| **B5 Surface Area Percentage** | 2.56568014863354e-06 |
| **B6 Surface Area Percentage** | 1.49045317165897e-54 |
| **B4 Band Power** | 9.37297391117984e-19 |
| **B5 Band Power** | 1.29793857874936e-29 |
| **B6 Band Power** | 8.65044513250506e-51 |
| **B4 Band Relative Power** | 3.50741965675334e-23 |
| **B5 Band Relative Power** | 2.83867664370904e-09 |
| **B6 Band Relative Power** | 3.05502393353002e-63 |

**Supp. Table 2.** This table presents the outcomes of the model selection process, showing the average log scores achieved by each model on the test dataset across 20 bootstrap iterations using harmonized data. The evaluation covers each spectral feature across five models of varying complexity: Model A (linear), Model B (smooth mean), Model C (smooth mean and variance), Model D (SHASH with fixed Kurtosis and Skewness), and Model E (full SHASH).

| Feature | Model A | Model B | Model C | Model D | Model E | Best Model |
| --- | --- | --- | --- | --- | --- | --- |
| **B4 Band Power** | -4.5917 | -4.2902 | -4.0268 | **-4.0210** | -4.0569 | Model D |
| **B4 Band Relative Power** | 0.0180 | 0.5499 | 0.5486 | 0.5862 | **0.5924** | Model E |
| **B4 Surface Area Percentage** | 0.5637 | 1.1629 | 1.1667 | **1.1776** | 1.1676 | Model D |
| **B5 Band Power** | -5.6355 | -5.0695 | -4.4595 | -4.3893 | **-4.3824** | Model E |
| **B5 Band Relative Power** | 0.3845 | 0.6087 | 0.6142 | 0.6238 | **0.6304** | Model E |
| **B5 Surface Area Percentage** | 1.1333 | 1.1679 | **1.1816** | 1.1800 | 1.1766 | Model C |
| **B6 Band Power** | -5.2543 | -4.6879 | -3.6071 | -3.3411 | **-3.3353** | Model E |
| **B6 Band Relative Power** | -0.0417 | -0.0074 | 0.0657 | 0.0967 | **0.1035** | Model E |
| **B6 Surface Area Percentage** | 0.5588 | 0.6510 | 0.6791 | 0.6827 | **0.6918** | Model E |

## References

1. Akula, S.K., Exposito-Alonso, D., Walsh, C.A., 2023. Shaping the brain: The emergence of cortical structure and folding. Dev. Cell 58, 2836–2849. 10.1016/j.devcel.2023.11.004

2. Bartha-Doering, L., Kollndorfer, K., Schwartz, E., Fischmeister, F.Ph.S., Langs, G., Weber, M., Lackner-Schmelz, S., Kienast, P., Stümpflen, M., Taymourtash, A., Mandl, S., Alexopoulos, J., Prayer, D., Seidl, R., Kasprian, G., 2023. Fetal temporal sulcus depth asymmetry has prognostic value for language development. Commun. Biol. 6, 109. 10.1038/s42003-023-04503-z

3. Bayly, P.V., Taber, L.A., Kroenke, C.D., 2014. Mechanical forces in cerebral cortical folding: A review of measurements and models. J. Mech. Behav. Biomed. Mater. 29, 568–581. 10.1016/j.jmbbm.2013.02.018

4. Bethlehem, R.A.I., Seidlitz, J., White, S.R., Vogel, J.W., Anderson, K.M., Adamson, C., Adler, S., Alexopoulos, G.S., Anagnostou, E., Areces-Gonzalez, A., Astle, D.E., Auyeung, B., Ayub, M., Bae, J., Ball, G., Baron-Cohen, S., Beare, R., Bedford, S.A., Benegal, V., Beyer, F., Blangero, J., Blesa Cábez, M., Boardman, J.P., Borzage, M., Bosch-Bayard, J.F., Bourke, N., Calhoun, V.D., Chakravarty, M.M., Chen, C., Chertavian, C., Chetelat, G., Chong, Y.S., Cole, J.H., Corvin, A., Costantino, M., Courchesne, E., Crivello, F., Cropley, V.L., Crosbie, J., Crossley, N., Delarue, M., Delorme, R., Desrivieres, S., Devenyi, G.A., Di Biase, M.A., Dolan, R., Donald, K.A., Donohoe, G., Dunlop, K., Edwards, A.D., Elison, J.T., Ellis, C.T., Elman, J.A., Eyler, L., Fair, D.A., Feczko, E., Fletcher, P.C., Fonagy, P., Franz, C.E., Galan-Garcia, L., Gholipour, A., Giedd, J., Gilmore, J.H., Glahn, D.C., Goodyer, I.M., Grant, P.E., Groenewold, N.A., Gunning, F.M., Gur, R.E., Gur, R.C., Hammill, C.F., Hansson, O., Hedden, T., Heinz, A., Henson, R.N., Heuer, K., Hoare, J., Holla, B., Holmes, A.J., Holt, R., Huang, H., Im, K., Ipser, J., Jack, C.R., Jackowski, A.P., Jia, T., Johnson, K.A., Jones, P.B., Jones, D.T., Kahn, R.S., Karlsson, H., Karlsson, L., Kawashima, R., Kelley, E.A., Kern, S., Kim, K.W., Kitzbichler, M.G., Kremen, W.S., Lalonde, F., Landeau, B., Lee, S., Lerch, J., Lewis, J.D., Li, J., Liao, W., Liston, C., Lombardo, M.V., Lv, J., Lynch, C., Mallard, T.T., Marcelis, M., Markello, R.D., Mathias, S.R., Mazoyer, B., McGuire, P., Meaney, M.J., Mechelli, A., Medic, N., Misic, B., Morgan, S.E., Mothersill, D., Nigg, J., Ong, M.Q.W., Ortinau, C., Ossenkoppele, R., Ouyang, M., Palaniyappan, L., Paly, L., Pan, P.M., Pantelis, C., Park, M.M., Paus, T., Pausova, Z., Paz-Linares, D., Pichet Binette, A., Pierce, K., Qian, X., Qiu, J., Qiu, A., Raznahan, A., Rittman, T., Rodrigue, A., Rollins, C.K., Romero-Garcia, R., Ronan, L., Rosenberg, M.D., Rowitch, D.H., Salum, G.A., Satterthwaite, T.D., Schaare, H.L., Schachar, R.J., Schultz, A.P., Schumann, G., Schöll, M., Sharp, D., Shinohara, R.T., Skoog, I., Smyser, C.D., Sperling, R.A., Stein, D.J., Stolicyn, A., Suckling, J., Sullivan, G., Taki, Y., Thyreau, B., Toro, R., Traut, N., Tsvetanov, K.A., Turk-Browne, N.B., Tuulari, J.J., Tzourio, C., Vachon-Presseau, É., Valdes-Sosa, M.J., Valdes-Sosa, P.A., Valk, S.L., Van Amelsvoort, T., Vandekar, S.N., Vasung, L., Victoria, L.W., Villeneuve, S., Villringer, A., Vértes, P.E., Wagstyl, K., Wang, Y.S., Warfield, S.K., Warrier, V., Westman, E., Westwater, M.L., Whalley, H.C., Witte, A.V., Yang, N., Yeo, B., Yun, H., Zalesky, A., Zar, H.J., Zettergren, A., Zhou, J.H., Ziauddeen, H., Zugman, A., Zuo, X.N., 3R-BRAIN, AIBL, Rowe, C., Alzheimer’s Disease Neuroimaging Initiative, Alzheimer’s Disease Repository Without Borders Investigators, Frisoni, G.B., CALM Team, Cam-CAN, CCNP, COBRE, cVEDA, ENIGMA Developmental Brain Age Working Group, Developing Human Connectome Project, FinnBrain, Harvard Aging Brain Study, IMAGEN, KNE96, The Mayo Clinic Study of Aging, NSPN, POND, The PREVENT-AD Research Group, Binette, A.P., VETSA, Bullmore, E.T., Alexander-Bloch, A.F., 2022. Brain charts for the human lifespan. Nature 604, 525–533. 10.1038/s41586-022-04554-y

5. Brossard-Racine, M., Du Plessis, A., Vezina, G., Robertson, R., Donofrio, M., Tworetzky, W., Limperopoulos, C., 2016. Brain Injury in Neonates with Complex Congenital Heart Disease: What Is the Predictive Value of MRI in the Fetal Period? Am. J. Neuroradiol. 37, 1338–1346. 10.3174/ajnr.A4716

6. Chi, J.G., Dooling, E.C., Gilles, F.H., 1977. Gyral development of the human brain. Ann. Neurol. 1, 86–93. 10.1002/ana.410010109

7. Ciceri, T., Casartelli, L., Montano, F., Conte, S., Squarcina, L., Bertoldo, A., Agarwal, N., Brambilla, P., Peruzzo, D., 2024. Fetal brain MRI atlases and datasets: A review. NeuroImage 292, 120603. 10.1016/j.neuroimage.2024.120603

8. Clouchoux, C., Kudelski, D., Gholipour, A., Warfield, S.K., Viseur, S., Bouyssi-Kobar, M., Mari, J.-L., Evans, A.C., Du Plessis, A.J., Limperopoulos, C., 2012. Quantitative in vivo MRI measurement of cortical development in the fetus. Brain Struct. Funct. 217, 127–139. 10.1007/s00429-011-0325-x

9. De Vareilles, H., Rivière, D., Mangin, J., Dubois, J., 2023. Development of cortical folds in the human brain: An attempt to review biological hypotheses, early neuroimaging investigations and functional correlates. Dev. Cogn. Neurosci. 61, 101249. 10.1016/j.dcn.2023.101249

10. Demirci, N., Holland, M.A., 2022. Cortical thickness systematically varies with curvature and depth in healthy human brains. Hum. Brain Mapp. 43, 2064–2084. 10.1002/hbm.25776

11. Dinga, R., Fraza, C.J., Bayer, J.M.M., Kia, S.M., Beckmann, C.F., Marquand, A.F., 2021. Normative modeling of neuroimaging data using generalized additive models of location scale and shape. 10.1101/2021.06.14.448106

12. Dubois, J., Benders, M., Borradori-Tolsa, C., Cachia, A., Lazeyras, F., Ha-Vinh Leuchter, R., Sizonenko, S.V., Warfield, S.K., Mangin, J.F., Hüppi, P.S., 2008. Primary cortical folding in the human newborn: an early marker of later functional development. Brain 131, 2028–2041. 10.1093/brain/awn137

13. Dubois, J., Lefèvre, J., Angleys, H., Leroy, F., Fischer, C., Lebenberg, J., Dehaene-Lambertz, G., Borradori-Tolsa, C., Lazeyras, F., Hertz-Pannier, L., Mangin, J.-F., Hüppi, P.S., Germanaud, D., 2019. The dynamics of cortical folding waves and prematurity-related deviations revealed by spatial and spectral analysis of gyrification. NeuroImage 185, 934–946. 10.1016/j.neuroimage.2018.03.005

14. Economo, C. von, Koskinas, G.N., Triárchou, L.K., 2008. Atlas of cytoarchitectonics of the adult human cerebral cortex: compiled at the Psychiatric Clinic of Hofrat Julius Wagner von Jauregg, Vienna. Karger, Basel.

15. Egaña-Ugrinovic, G., Sanz-Cortes, M., Figueras, F., Bargalló, N., Gratacós, E., 2013. Differences in cortical development assessed by fetal MRI in late-onset intrauterine growth restriction. Am. J. Obstet. Gynecol. 209, 126.e1–126.e8. 10.1016/j.ajog.2013.04.008

16. Feess-Higgins, A., Larroche, J.-C., 1987. Le Développement du cerveau fœtal humain: atlas anatomique. INSERM Masson, Paris.

17. Garcia, K.E., Kroenke, C.D., Bayly, P.V., 2018. Mechanics of cortical folding: stress, growth and stability. Philos. Trans. R. Soc. B Biol. Sci. 373, 20170321. 10.1098/rstb.2017.0321

18. Germanaud, D., Lefèvre, J., Fischer, C., Bintner, M., Curie, A., Des Portes, V., Eliez, S., Elmaleh-Bergès, M., Lamblin, D., Passemard, S., Operto, G., Schaer, M., Verloes, A., Toro, R., Mangin, J.F., Hertz-Pannier, L., 2014. Simplified gyral pattern in severe developmental microcephalies? New insights from allometric modeling for spatial and spectral analysis of gyrification. NeuroImage 102, 317–331. 10.1016/j.neuroimage.2014.07.057

19. Germanaud, D., Lefèvre, J., Toro, R., Fischer, C., Dubois, J., Hertz-Pannier, L., Mangin, J.-F., 2012. Larger is twistier: Spectral analysis of gyrification (SPANGY) applied to adult brain size polymorphism. NeuroImage 63, 1257–1272. 10.1016/j.neuroimage.2012.07.053

20. Gholipour, A., Rollins, C.K., Velasco-Annis, C., Ouaalam, A., Akhondi-Asl, A., Afacan, O., Ortinau, C.M., Clancy, S., Limperopoulos, C., Yang, E., Estroff, J.A., Warfield, S.K., 2017. A normative spatiotemporal MRI atlas of the fetal brain for automatic segmentation and analysis of early brain growth. Sci. Rep. 7, 476. 10.1038/s41598-017-00525-w

21. Im, K., Grant, P.E., 2019. Sulcal pits and patterns in developing human brains. NeuroImage 185, 881–890. 10.1016/j.neuroimage.2018.03.057

22. Ji, L., Menu, I., Majbri, A., Bhatia, T., Trentacosta, C.J., Thomason, M.E., 2024. Trajectories of human brain functional connectome maturation across the birth transition. PLOS Biol. 22, e3002909. 10.1371/journal.pbio.3002909

23. Juergensen, S., Liu, J., Xu, D., Zhao, Y., Moon Grady, A.J., Glenn, O., McQuillen, P., Peyvandi, S., 2024. Fetal circulatory physiology and brain development in complex congenital heart disease: A multi modal imaging study. Prenat. Diagn. 44, 856–864. 10.1002/pd.6450

24. Kippenhan, J.S., Olsen, R.K., Mervis, C.B., Morris, C.A., Kohn, P., Meyer-Lindenberg, A., Berman, K.F., 2005. Genetic Contributions to Human Gyrification: Sulcal Morphometry in Williams Syndrome. J. Neurosci. 25, 7840–7846. 10.1523/JNEUROSCI.1722-05.2005

25. Kostović, I., Sedmak, G., Judaš, M., 2019. Neural histology and neurogenesis of the human fetal and infant brain. NeuroImage 188, 743–773. 10.1016/j.neuroimage.2018.12.043

26. Lamballais, S., Vinke, E.J., Vernooij, M.W., Ikram, M.A., Muetzel, R.L., 2020. Cortical gyrification in relation to age and cognition in older adults. NeuroImage 212, 116637. 10.1016/j.neuroimage.2020.116637

27. Lefèvre, J., Mangin, J.-F., 2010. A Reaction-Diffusion Model of Human Brain Development. PLoS Comput. Biol. 6, e1000749. 10.1371/journal.pcbi.1000749

28. Lefèvre, J., Pepe, A., Muscato, J., De Guio, F., Girard, N., Auzias, G., Germanaud, D., 2018. SPANOL (SPectral ANalysis of Lobes): A Spectral Clustering Framework for Individual and Group Parcellation of Cortical Surfaces in Lobes. Front. Neurosci. 12, 354. 10.3389/fnins.2018.00354

29. Llinares-Benadero, C., Borrell, V., 2019. Deconstructing cortical folding: genetic, cellular and mechanical determinants. Nat. Rev. Neurosci. 20, 161–176. 10.1038/s41583-018-0112-2

30. Marquand, A.F., Rezek, I., Buitelaar, J., Beckmann, C.F., 2016. Understanding Heterogeneity in Clinical Cohorts Using Normative Models: Beyond Case-Control Studies. Biol. Psychiatry 80, 552–561. 10.1016/j.biopsych.2015.12.023

31. Mihailov, A., Pron, A., Lefèvre, J., Deruelle, C., Desnous, B., Bretelle, F., Manchon, A., Milh, M., Rousseau, F., Girard, N., Auzias, G., 2025. Burst of gyrification in the human brain after birth. Commun. Biol. 8, 805. 10.1038/s42003-025-08155-z

32. Pomponio, R., Erus, G., Habes, M., Doshi, J., Srinivasan, D., Mamourian, E., Bashyam, V., Nasrallah, I.M., Satterthwaite, T.D., Fan, Y., Launer, L.J., Masters, C.L., Maruff, P., Zhuo, C., Völzke, H., Johnson, S.C., Fripp, J., Koutsouleris, N., Wolf, D.H., Gur, Raquel, Gur, Ruben, Morris, J., Albert, M.S., Grabe, H.J., Resnick, S.M., Bryan, R.N., Wolk, D.A., Shinohara, R.T., Shou, H., Davatzikos, C., 2020. Harmonization of large MRI datasets for the analysis of brain imaging patterns throughout the lifespan. NeuroImage 208, 116450. 10.1016/j.neuroimage.2019.116450

33. Rabiei, H., Richard, F., Coulon, O., Lefevre, J., 2017. Local Spectral Analysis of the Cerebral Cortex: New Gyrification Indices. IEEE Trans. Med. Imaging 36, 838–848. 10.1109/TMI.2016.2633393

34. Rajagopalan, V., Scott, J., Habas, P.A., Kim, K., Corbett-Detig, J., Rousseau, F., Barkovich, A.J., Glenn, O.A., Studholme, C., 2011. Local Tissue Growth Patterns Underlying Normal Fetal Human Brain Gyrification Quantified *In Utero*. J. Neurosci. 31, 2878–2887. 10.1523/JNEUROSCI.5458-10.2011

35. Robinson, E.C., Garcia, K., Glasser, M.F., Chen, Z., Coalson, T.S., Makropoulos, A., Bozek, J., Wright, R., Schuh, A., Webster, M., Hutter, J., Price, A., Cordero Grande, L., Hughes, E., Tusor, N., Bayly, P.V., Van Essen, D.C., Smith, S.M., Edwards, A.D., Hajnal, J., Jenkinson, M., Glocker, B., Rueckert, D., 2018. Multimodal surface matching with higher-order smoothness constraints. NeuroImage 167, 453–465. 10.1016/j.neuroimage.2017.10.037

36. Rutherford, S., Barkema, P., Tso, I.F., Sripada, C., Beckmann, C.F., Ruhe, H.G., Marquand, A.F., 2023. Evidence for embracing normative modeling. eLife 12, e85082. 10.7554/eLife.85082

37. Sanchez, T., Martí-Juan, G., Meunier, D., Ballester, M.A.G., Camara, O., Piella, G., Cuadra, M.B., Auzias, G., 2025. Fetpype: An Open-Source Pipeline for Reproducible Fetal Brain MRI Analysis. 10.48550/arXiv.2512.17472

38. Sanchez, T., Mihailov, A., Martí-Juan, G., Girard, N., Manchon, A., Milh, M., Eixarch, E., Dunet, V., Koob, M., Pomar, L., Sichitiu, J., Ballester, M.A.G., Camara, O., Piella, G., Cuadra, M.B., Auzias, G., n.d. Data quality biases normative models derived from fetal brain MRI.

39. Schwartz, E., Diogo, M.C., Glatter, S., Seidl, R., Brugger, P.C., Gruber, G.M., Kiss, H., Nenning, K.-H., IRC5 consortium, Langs, G., Prayer, D., Kasprian, G., 2021. The Prenatal Morphomechanic Impact of Agenesis of the Corpus Callosum on Human Brain Structure and Asymmetry. Cereb. Cortex bhab066. 10.1093/cercor/bhab066

40. Solhtalab, A., Guo, Y., Gholipour, A., Dai, W., Razavi, M.J., 2025. Mechanics of the Spatiotemporal Evolution of Sulcal Pits in the Folding Brain. Hum. Brain Mapp. 46, e70332. 10.1002/hbm.70332

41. Stasinopoulos, M., Rigby, R., 2012. gamlss: Generalized Additive Models for Location Scale and Shape. 10.32614/CRAN.package.gamlss

42. Tallinen, T., Chung, J.Y., Rousseau, F., Girard, N., Lefèvre, J., Mahadevan, L., 2016. On the growth and form of cortical convolutions. Nat. Phys. 12, 588–593. 10.1038/nphys3632

43. Thompson, P.M., Lee, A.D., Dutton, R.A., Geaga, J.A., Hayashi, K.M., Eckert, M.A., Bellugi, U., Galaburda, A.M., Korenberg, J.R., Mills, D.L., Toga, A.W., Reiss, A.L., 2005. Abnormal Cortical Complexity and Thickness Profiles Mapped in Williams Syndrome. J. Neurosci. 25, 4146–4158. 10.1523/JNEUROSCI.0165-05.2005

44. Tustison, N.J., Cook, P.A., Holbrook, A.J., Johnson, H.J., Muschelli, J., Devenyi, G.A., Duda, J.T., Das, S.R., Cullen, N.C., Gillen, D.L., Yassa, M.A., Stone, J.R., Gee, J.C., Avants, B.B., 2021. The ANTsX ecosystem for quantitative biological and medical imaging. Sci. Rep. 11, 9068. 10.1038/s41598-021-87564-6

45. Uus, A.U., Kyriakopoulou, V., Makropoulos, A., Fukami-Gartner, A., Cromb, D., Davidson, A., Cordero-Grande, L., Price, A.N., Grigorescu, I., Williams, L.Z.J., Robinson, E.C., Lloyd, D., Pushparajah, K., Story, L., Hutter, J., Counsell, S.J., Edwards, A.D., Rutherford, M.A., Hajnal, J.V., Deprez, M., 2023. BOUNTI: Brain vOlumetry and aUtomated parcellatioN for 3D feTal MRI. 10.7554/eLife.88818.1

46. Wagstyl, K., Lepage, C., Bludau, S., Zilles, K., Fletcher, P.C., Amunts, K., Evans, A.C., 2018. Mapping Cortical Laminar Structure in the 3D BigBrain. Cereb. Cortex 28, 2551–2562. 10.1093/cercor/bhy074

47. Walhovd, K.B., Krogsrud, S.K., Amlien, I.K., Sørensen, Ø., Wang, Y., Bråthen, A.C.S., Overbye, K., Kransberg, J., Mowinckel, A.M., Magnussen, F., Herud, M., Håberg, A.K., Fjell, A.M., Vidal-Pineiro, D., 2024. Fetal influence on the human brain through the lifespan. eLife 12, RP86812. 10.7554/eLife.86812

48. Wang, X., Lefèvre, J., Bohi, A., Harrach, M.A., Dinomais, M., Rousseau, F., 2021. The influence of biophysical parameters in a biomechanical model of cortical folding patterns. Sci. Rep. 11, 7686. 10.1038/s41598-021-87124-y

49. Wang, X., Studholme, C., Grigsby, P.L., Frias, A.E., Cuzon Carlson, V.C., Kroenke, C.D., 2017. Folding, But Not Surface Area Expansion, Is Associated with Cellular Morphological Maturation in the Fetal Cerebral Cortex. J. Neurosci. 37, 1971–1983. 10.1523/JNEUROSCI.3157-16.2017

50. Wu, Y., Lu, Y.-C., Jacobs, M., Pradhan, S., Kapse, K., Zhao, L., Niforatos-Andescavage, N., Vezina, G., Du Plessis, A.J., Limperopoulos, C., 2020. Association of Prenatal Maternal Psychological Distress With Fetal Brain Growth, Metabolism, and Cortical Maturation. JAMA Netw. Open 3, e1919940. 10.1001/jamanetworkopen.2019.19940

51. Xu, J., Moyer, D., Gagoski, B., Iglesias, J.E., Grant, P.E., Golland, P., Adalsteinsson, E., 2023. NeSVoR: Implicit Neural Representation for Slice-to-Volume Reconstruction in MRI. IEEE Trans. Med. Imaging 42, 1707–1719. 10.1109/TMI.2023.3236216

52. Xu, X., Sun, C., Sun, J., Shi, W., Shen, Y., Zhao, R., Luo, W., Li, M., Wang, G., Wu, D., 2022. Spatiotemporal Atlas of the Fetal Brain Depicts Cortical Developmental Gradient. J. Neurosci. 42, 9435–9449. 10.1523/JNEUROSCI.1285-22.2022

53. Yehuda, B., Rabinowich, A., Link-Sourani, D., Avisdris, N., Ben-Zvi, O., Specktor-Fadida, B., Joskowicz, L., Ben-Sira, L., Miller, E., Ben Bashat, D., 2023. Automatic Quantification of Normal Brain Gyrification Patterns and Changes in Fetuses with Polymicrogyria and Lissencephaly Based on MRI. Am. J. Neuroradiol. 44, 1432–1439. 10.3174/ajnr.A8046

54. Yin, S., Liu, C., Choi, G.P., Jung, Y., Heuer, K., Toro, R., Mahadevan, L., 2025. Morphogenesis and morphometry of brain folding patterns across species. eLife 14, RP107138. 10.7554/eLife.107138.3

55. Yun, H.J., Vasung, L., Tarui, T., Rollins, C.K., Ortinau, C.M., Grant, P.E., Im, K., 2020. Temporal Patterns of Emergence and Spatial Distribution of Sulcal Pits During Fetal Life. Cereb. Cortex 30, 4257–4268. 10.1093/cercor/bhaa053

