## Supplementary figures and images for "Characterization of Fetal Cortical Development Using Spectral Analysis of Gyrification (SPANGY)"

### Supp. Fig. 1

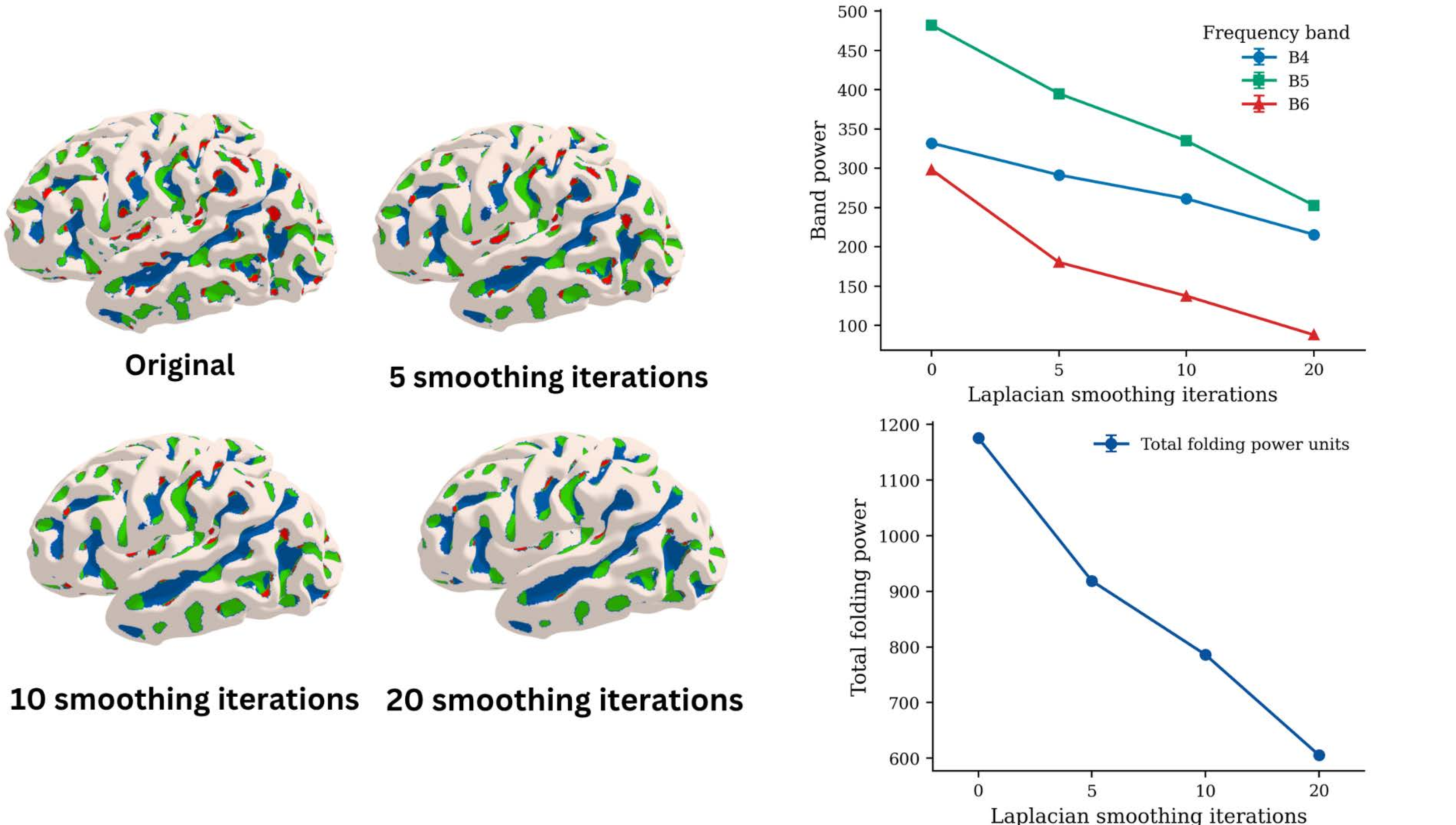

### Supp. Fig. 2

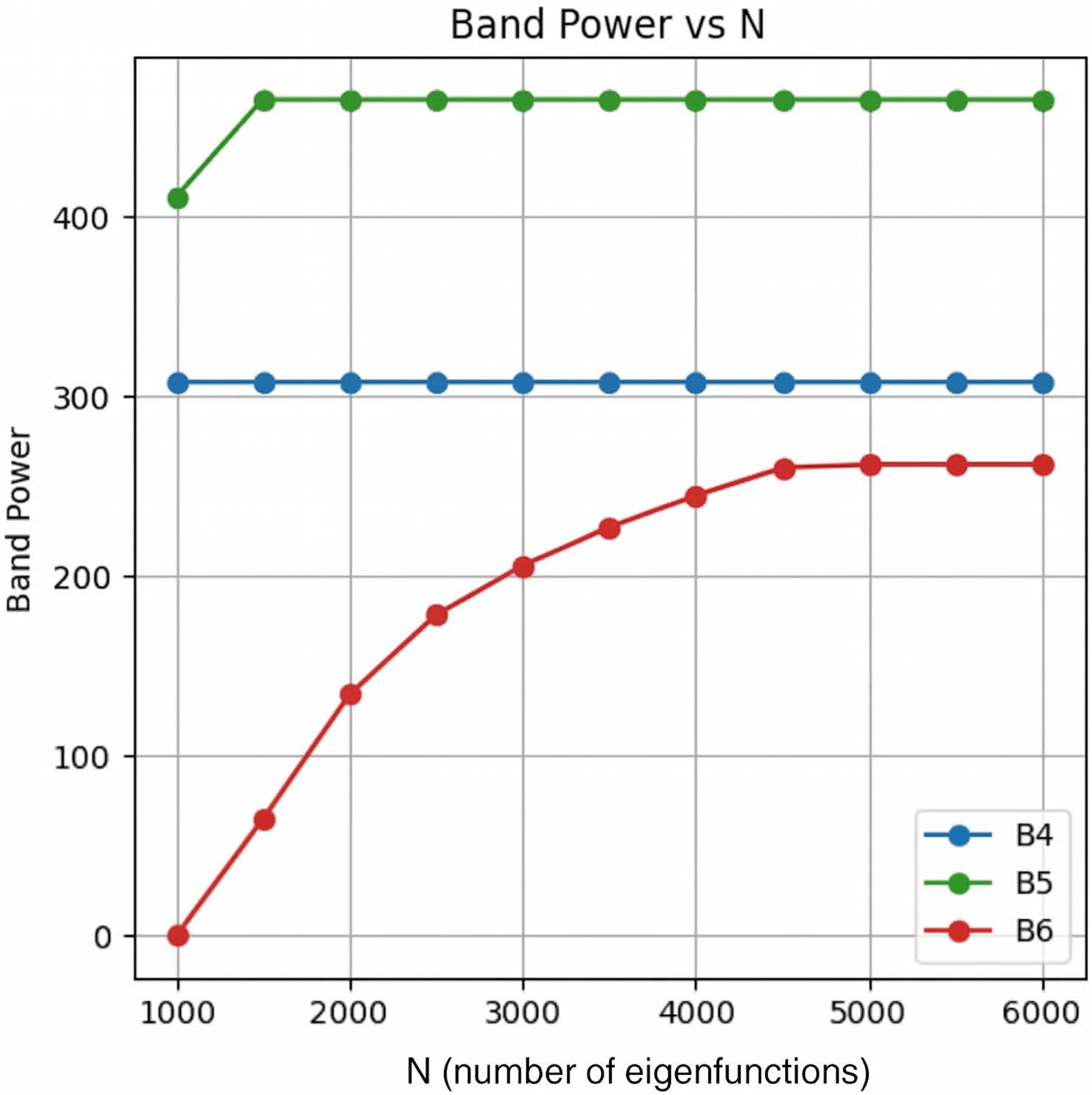

### Supp. FIg. 3

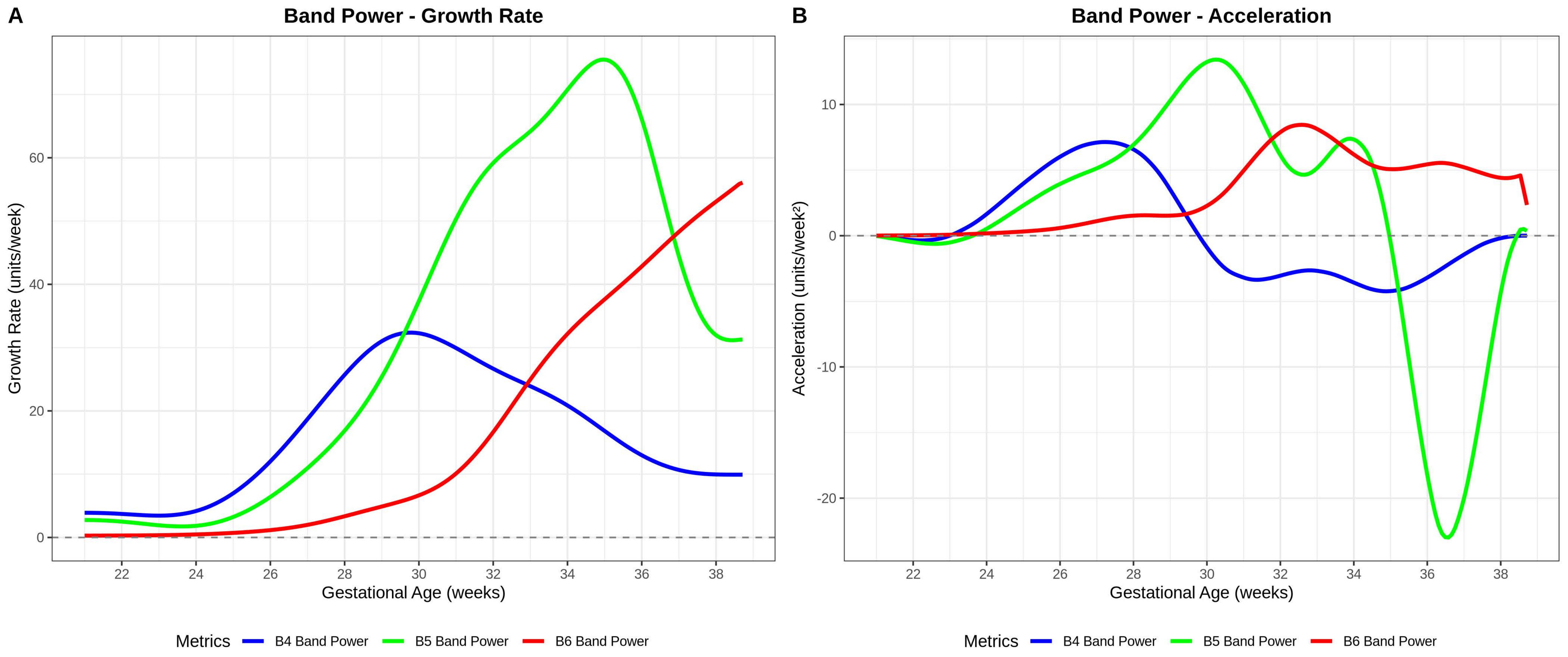

### Supp. Fig. 4

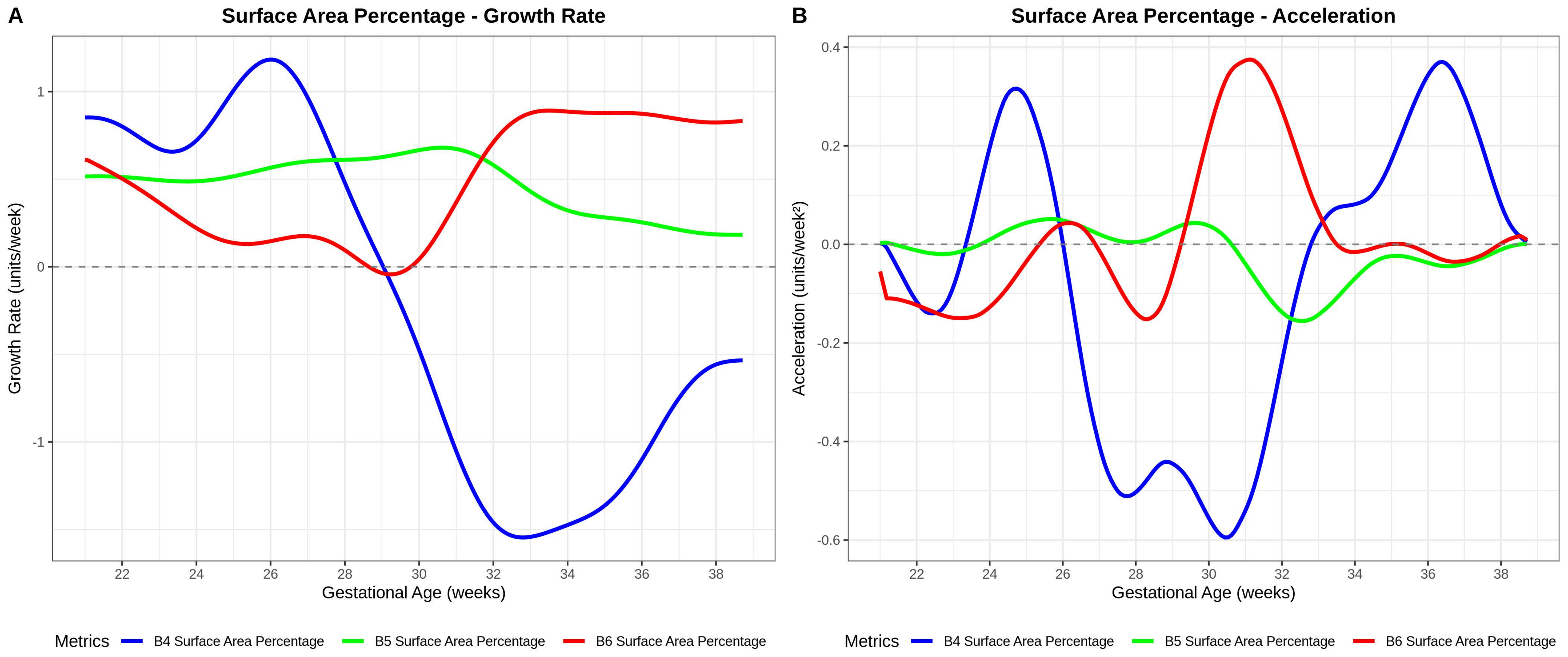

### Supp. Fig. 5

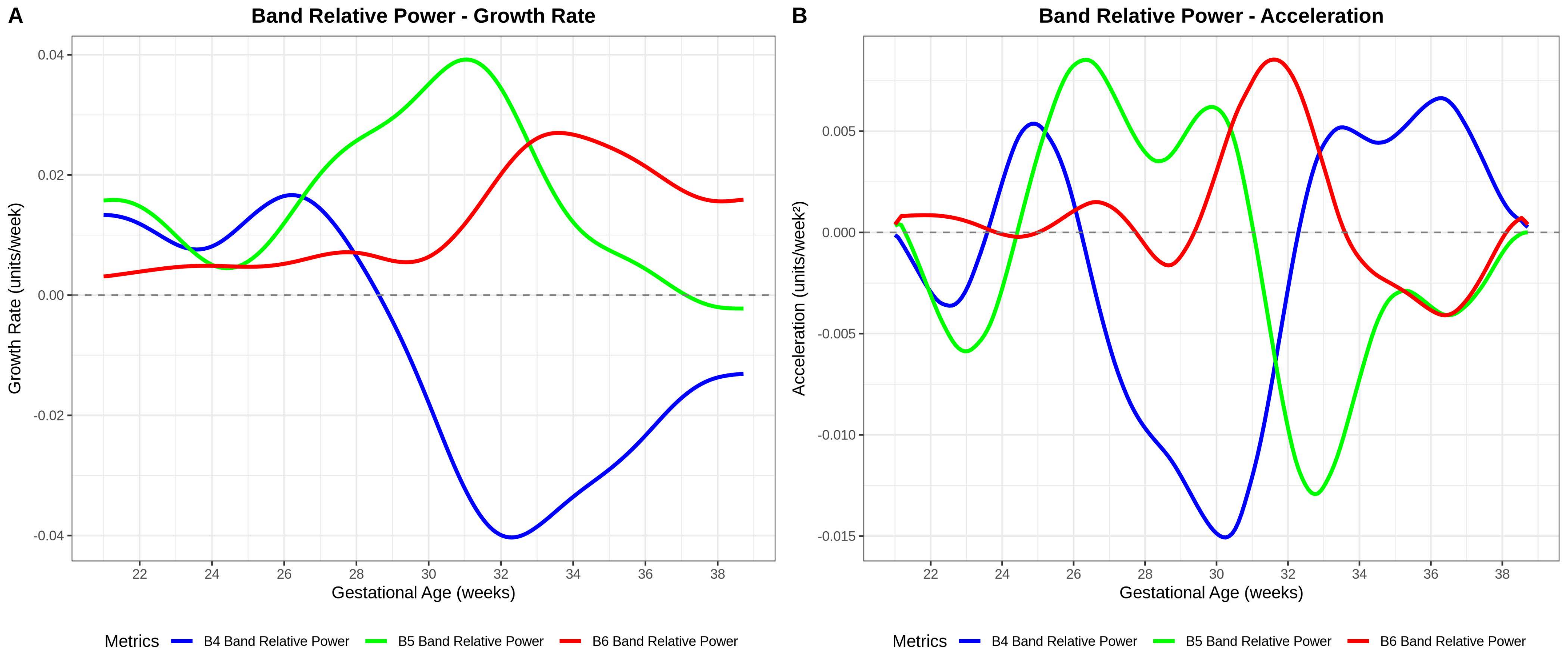

### supp. Fig. 7

#### Relationship between the absolute band powers for B4-B6 and smoothing across different ages.

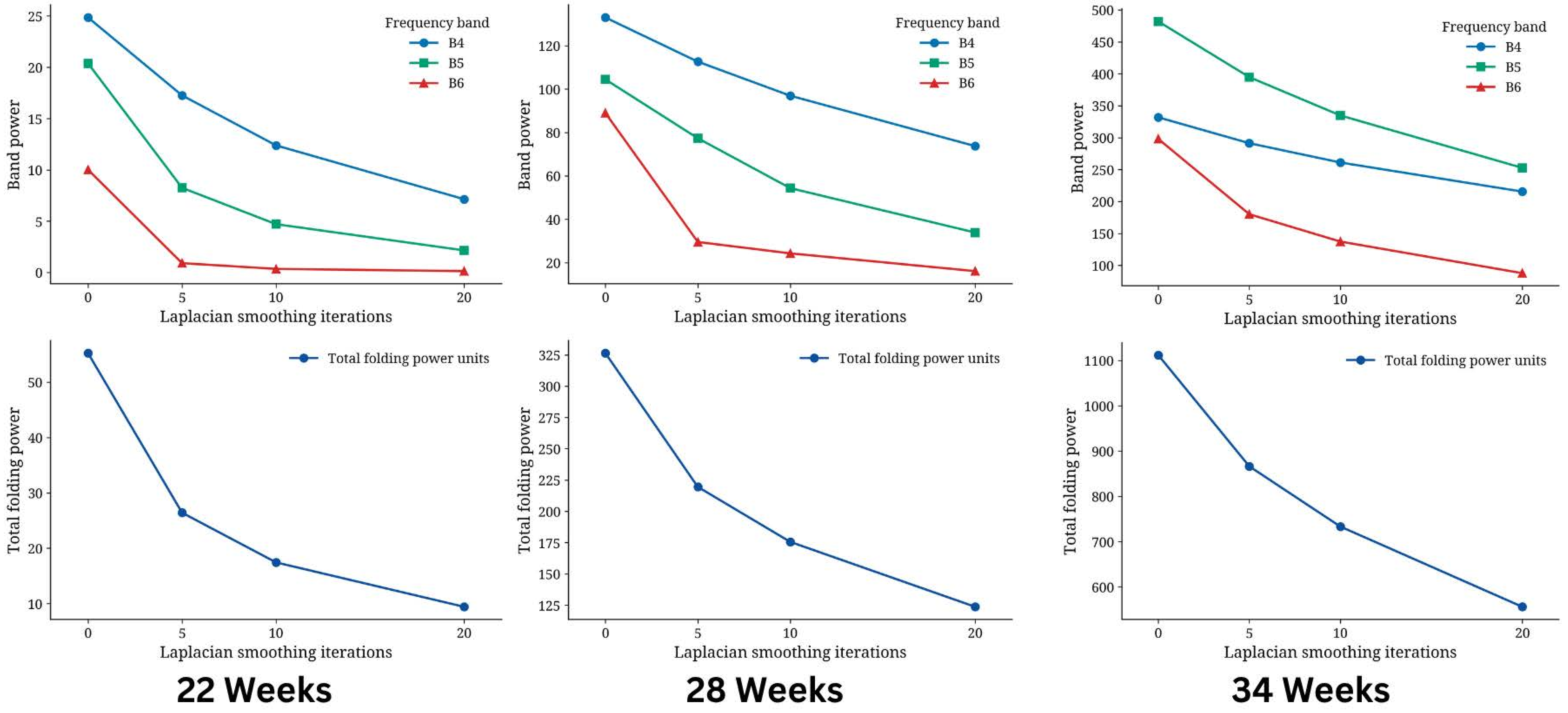
