## Supplementary material for "Characterization of Fetal Cortical Development Using Spectral Analysis of Gyrification (SPANGY)": Supp. Fig. 6

### Relationship between the number of eigenfunctions (N) and the absolute band powers for B4-B6 across different ages.

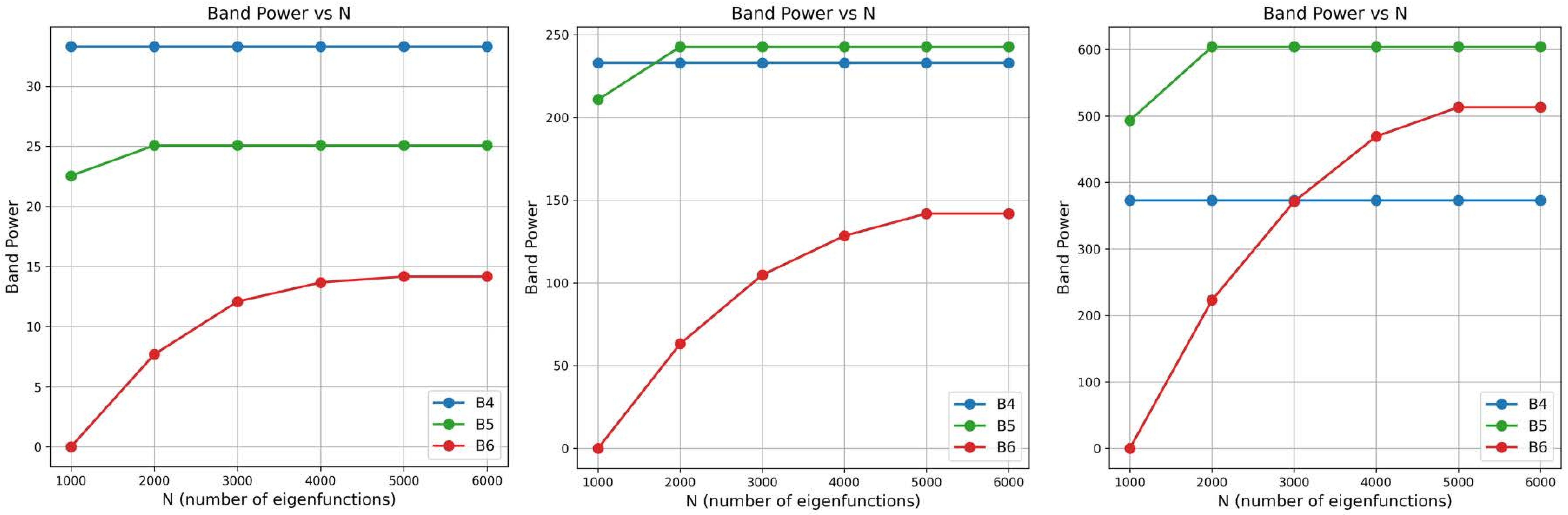


**24 Weeks** **32 Weeks** **37 Weeks**
